# Rac1 and CHK1 Converge on Abi-1 to Regulate DNA Repair Dependency and Treatment Response in Human Cancers

**DOI:** 10.64898/2026.09.11.750952

**Authors:** Yiheng Huang, Elizabeth McCulla, Jihan Park, Annabel Yang, Timothy Jho, Angelica Lin, Jie Xu, Kari Wilder-Romans, Luke D. Hess, Jing Li, Ningning Liang, Ziqing Zhu, Deven Kothari, Karen Jin, Sarah Kim, Sanduni H. Premathilaka, Bo Wen, Duxin Sun, Michelle Vinco, Sean P. Ferris, Meredith A. Morgan, Theodore S. Lawrence, Yatrik M. Shah, Menggang Yu, Molly E. Heft Neal, J. Chad Brenner, Suranganie Dharmawardhane, Jose F. Rodriguez-Orengo, Daniel R. Wahl, Weihua Zhou

## Abstract

Radiation therapy (RT) resistance remains a major clinical challenge, yet biomarkers guiding precision radiosensitization are lacking. We previously demonstrated that Rac1 promotes RT resistance in glioblastoma (GBM) by inducing Abi-1-S323 dephosphorylation and enhancing non-homologous end joining (NHEJ). Here, we identify Abi-1-S323 as a key regulator of DNA repair states and a determinant of therapeutic efficacy in human cancers. Clinically, loss of Abi-1-S323 phosphorylation was associated with poor outcomes in patients with RT-treated GBM. Bioinformatic analyses revealed that non-small cell lung cancer (NSCLC) and head and neck cancer (HNC) are among the cancers with frequent *RAC1* amplification, suggesting that these tumor types may have increased Rac1-Abi-1 signaling activity. Loss of Abi-1-S323 phosphorylation also predicted poor outcomes in patients with RT-treated HNC. Consistent with these clinical observations, high Rac1 activity and low Abi-1-S323 phosphorylation were associated with enhanced DNA double-strand break repair and radioresistance in NSCLC and HNC models, whereas genetic or pharmacological inhibition of this signaling impaired DNA repair and radiosensitized tumors *in vitro* and *in vivo*. Mechanistically, we identified CHK1 as a kinase that phosphorylates Abi-1 at S323 and defines an alternative homologous recombination (HR)-dependent repair state. Tumors with high Rac1-Abi-1 signaling exhibited elevated NHEJ capacity and were selectively radiosensitized by Rac1 inhibition, whereas tumors with low Rac1-Abi-1 signaling displayed high CHK1 activity, preferentially relied on HR, and were selectively radiosensitized by CHK1 inhibition. These findings establish Abi-1-S323 as a biomarker defining therapeutically distinct DNA repair states and provide a framework for precision radiosensitization.

**Significance:** Radiation resistance remains a major barrier to cancer treatment. We identify Abi-1-S323 phosphorylation as a biomarker of distinct DNA repair states and therapeutic vulnerabilities, providing a framework for precision radiosensitization.

## Introduction

Genotoxic radiation therapy (RT) is a cornerstone of cancer treatment. However, RT resistance remains a major clinical challenge. RT exerts its therapeutic effects primarily by inducing DNA double-strand breaks (DSBs), the most cytotoxic form of DNA damage. Efficient repair of RT-induced DSBs, however, enables tumor cell survival, thereby limiting treatment efficacy and contributing to RT resistance and disease recurrence (1,2). The DNA damage response (DDR) is the principal cellular network that detects and repairs DSBs, predominantly through the homologous recombination (HR) and non-homologous end joining (NHEJ) pathways (3). Consequently, tumors with impaired DDR are generally more sensitive to RT (4), whereas those with enhanced DNA repair capacity exhibit intrinsic or acquired radioresistance (5,6). These observations have prompted extensive efforts to combine RT with inhibitors targeting the DDR, several of which have demonstrated promising activity in preclinical studies (7-11), and are currently being evaluated in clinical trials (12-14). Despite these advances, our understanding of the mechanisms that regulate DNA repair pathways and therapeutic response to RT in individual tumors remains incomplete. Understanding which DSB repair pathways predominate in a given tumor may uncover new opportunities for personalized DDR-directed radiosensitization strategies.

Metabolic signaling plays a critical role in regulating the DDR and genotoxic therapy responses (15-21). We recently identified a mechanistic link between GTP metabolism and RT resistance: increased GTP activates Rac1, which activates protein phosphatase 5 (PP5) and promotes dephosphorylation of serine 323 on Abl-interactor 1 (Abi-1), a protein previously not known to regulate DNA repair (22). Dephosphorylation of Abi-1-S323 promotes NHEJ and confers resistance to genotoxic RT in laboratory models of GBM. However, important questions remain regarding this biology. The kinase(s) responsible for mediating the inactivating phosphorylation of Abi-1-S323 are unknown, as is whether this phosphorylation influences the choice between HR and NHEJ for repairing RT-induced DSBs. Several important therapeutic questions also remain unanswered, including whether Abi-1 dephosphorylation occurs in human tumors and whether it is associated with RT resistance. Finally, it remains unknown whether this mechanism extends beyond gliomas, where it could represent a targetable regulator of RT resistance in different cancers.

Here, we demonstrate that Abi-1-S323 dephosphorylation predicts poor patient outcomes following RT and defines distinct DNA repair states and therapeutic vulnerabilities across human cancers. We first found that elevated Rac1-Abi-1 signaling, was associated with significantly poorer outcomes in patients with GBM, non-small cell lung cancer (NSCLC) and head and neck cancer (HNC). Consistent with these clinical observations, elevated Rac1 activity in NSCLC and HNC models was associated with reduced Abi-1-S323 phosphorylation, enhanced DSB repair, and intrinsic radioresistance. Genetic or pharmacological inhibition of Rac1 impaired DSB repair and radiosensitized Rac1-high, radioresistant cancer models *in vitro* and *in vivo*. Mechanistically, we identified CHK1 as a kinase that phosphorylates Abi-1 at S323, revealing distinct DNA repair dependencies associated with differential Abi-1 regulation. Tumors with high Rac1 activity and low Abi-1-S323 phosphorylation exhibited enhanced NHEJ and were selectively radiosensitized by Rac1 inhibition. In contrast, tumors with low Rac1 activity and high Abi-1-S323 phosphorylation displayed elevated CHK1 activity, preferentially relied on HR, and were selectively radiosensitized by CHK1 inhibition. These findings establish Abi-1-S323 phosphorylation as a biomarker of therapeutically distinct DNA repair states and provide a mechanistic foundation for biomarker-guided precision radiotherapy across human cancers.

## Material and Methods

### Patient Samples

A tumor microarray (TMA) of primary gliomas was constructed from formalin-fixed, paraffin-embedded (FFPE) blocks of pretreatment biopsy specimens collected by the University of Michigan Brain Tumor Bank (23). All tissue blocks were reviewed by a board-certified neuropathologist, who identified and marked representative tumor regions for TMA construction. TMAs were generated by the Tissue and Molecular Pathology Shared Resource (TMPSR) at the University of Michigan. Each block was represented by a 1mm-diameter core obtained from a representative area of the tumor, in addition to normal neuro samples and other normal tissues. The TMA originally included 74 patients, of whom 60 received RT following craniotomy biopsy and had available survival data. Among these 60 patients, 33 with World Health Organization (WHO) grade IV glioblastoma (GBM) were included in the survival analysis (**Fig. 1A** and **B**). The use of clinical specimens and associated clinical data was approved by the University of Michigan Medical School Institutional Review Board (IRBMED; HUM00165469).

**Figure 1.**
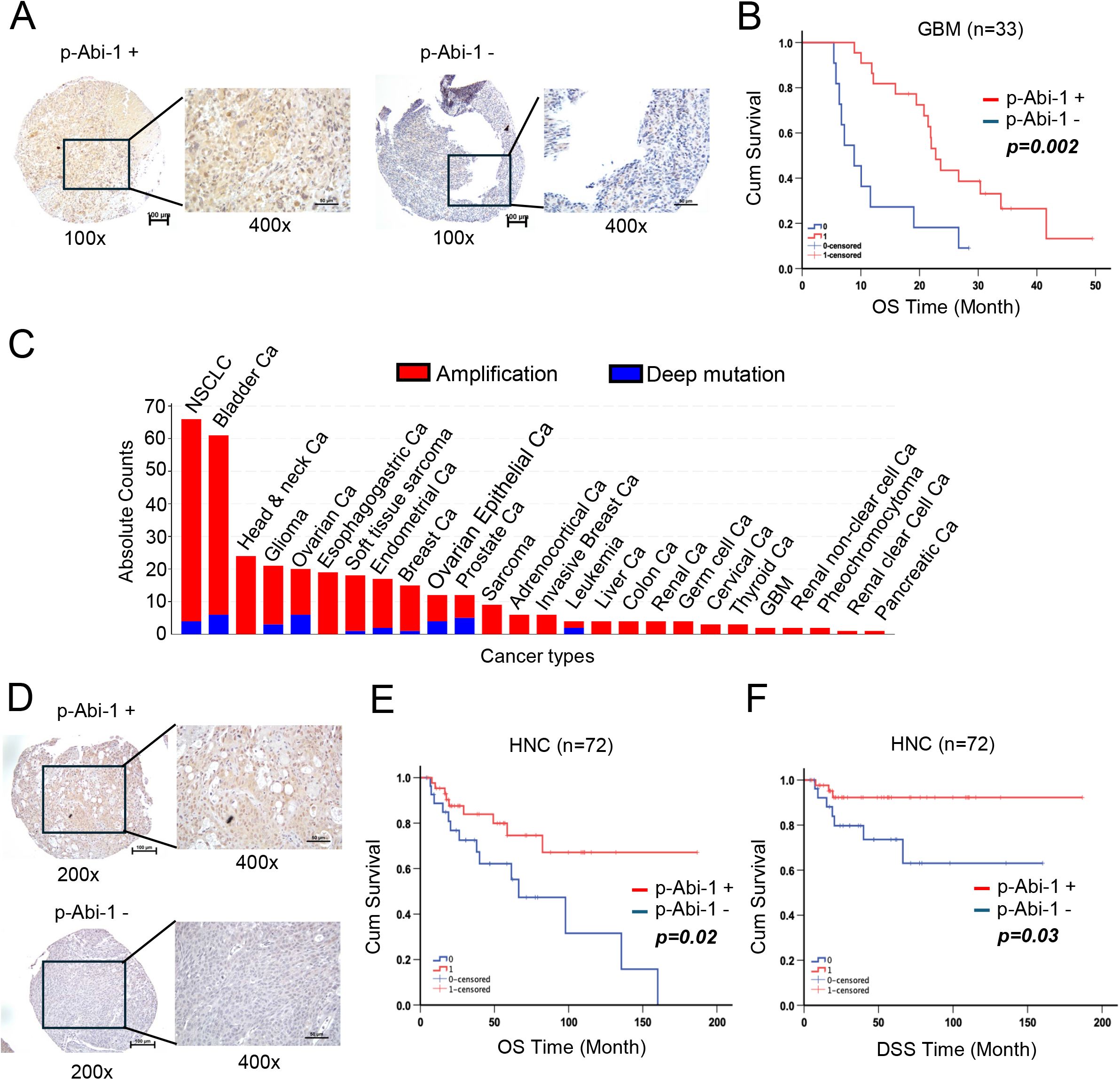
Abi-1-S323 phosphorylation predicts clinical outcome following RT. **(A)** p-Abi-1 S323 expression was assessed by IHC using a primary antibody at 1:300 dilution in a recently generated glioma TMA. Tumors were classified as p-Abi-1-S323-positive or -negative based on IHC staining. Representative images are shown. **(B)** Kaplan–Meier analysis of overall survival (OS) in GBM patients stratified by p-Abi-1-S323 expression. Median OS was 8.9 ± 1.9 months for patients with p-Abi-1-S323-negative tumors and 22.8 ± 1.3 months for those with p-Abi-1-S323-positive tumors (p = 0.002). **(C)** RNA-seq data from cBioPortal showing *RAC1* mRNA expression across cancer types. **(D)** p-Abi-1-S323 expression was assessed by IHC using a primary antibody at 1:200 dilution in an HNC TMA. A total of 72 HNC patients who received RT as part of their clinical care and had pre-treatment tumor specimens available were included in the analysis. Representative images of p-Abi-1-S323-positive and -negative tumors are shown. **(E-F)** Kaplan–Meier analyses of OS and disease-specific survival (DSS), respectively, in the 72 patients stratified by p-Abi-1 S323 expression. P values were calculated using the log-rank (Mantel–Cox) test for **(B), (E),** and **(F).**

TMA of primary laryngeal squamous cell carcinoma (LSCC) was previously constructed, and this study was approved by the University of Michigan (IRBMED; HUM00042189) (24,25). The TMAs initially included tumor specimens from 107 patients who underwent induction selection paradigms at our institution between 1995 and 2017, with specimens collected before induction therapy. Briefly, patients underwent 1-2 cycles of chemotherapy or chemotherapy combined with targeted therapy. Patients whose tumors did not improve by >50% proceeded to surgery with total laryngectomy, whereas patients whose tumors improved by >50% after induction therapy subsequently received definitive RT-based treatment. For patients who experienced tumor recurrence after definitive treatment, salvage surgery was performed followed by adjuvant RT. Of the 107 patients, 83 responded to the induction paradigm and subsequently received RT-based treatment and were included in this study. After excluding 11 patients whose tissue cores were lost or detached during TMA processing or staining, 72 patients were included in the survival analysis (**Fig. 1E** and **F**).

### *In vivo* xenograft models

Xenograft models were established as previously described (16,26). All animal experiments were approved by the University Committee on Use and Care of Animals (UCUCA) at the University of Michigan. Rag1 mice (6–8 weeks old; Strain# 002216) were obtained from The Jackson Laboratory and maintained under specific pathogen-free conditions. Animal housing was managed by the University of Michigan Unit for Laboratory Animal Medicine (ULAM), with a temperature of 74 °F, relative humidity of 30–70%, and a 12-hour light/12-hour dark cycle. For tumor implantation, cells were resuspended in a 1:1 mixture of PBS and Matrigel (BD Biosciences) and subcutaneously injected into the bilateral dorsal flanks of mice. When tumors reached approximately 100 mm³, mice were randomized into four treatment groups: vehicle control (0.5% (w/v) methylcellulose and 0.1% (v/v) polysorbate 80), MBQ-167 alone, RT alone, or the combination of MBQ-167 and RT. MBQ-167 (75 mg/kg) and RT (2 Gy/fraction) were administered according to the treatment schedules shown in Supplementary **Fig. S5A** and Supplementary **Fig. S5D**. A subset of tumors was harvested 2–4 hours after the second RT (or sham RT) dose for biological endpoint analyses, including mass spectrometry, pharmacokinetic (PK), immunoblotting, and immunohistochemistry. The remaining mice continued on their assigned treatment regimens, with tumor volume and body weight measured two to three times per week. Tumor volume was calculated from digital caliper measurements using the formula: (π/6) × Length × Width².

### Pharmacokinetic analyses of MBQ-167

Plasma and corresponding tumor samples were collected from mice in each treatment group as described in Supplementary Fig. S5A and S5D. MBQ-167 concentrations were quantified by the liquid chromatography–tandem mass spectrometry (LC–MS/MS) as described previously(27,28). Tumor tissues were homogenized in 20% acetonitrile, whereas plasma samples underwent protein precipitation with cold methanol before LC–MS/MS analysis. The LC−MS/MS method consisted of an Agilent 1200 series HPLC system and chromatographic separation of the tested compound was achieved using a Waters XBridge-C18 column (5 cm × 2.1 mm, 3.5 μm). An AB Sciex QTrap 3200 mass spectrometer equipped with an electrospray ionization source (Applied Biosystems, Toronto, Canada) in the positive-ion multiple reaction monitoring (MRM) mode was used for detection. The mobile phases were 0.1% formic acid in purified water (A) and 0.1% formic acid in acetonitrile (B). The gradient (B) was held at 10% (0-0.2 min), increased to 98% at 0.3 min, then stayed at isocratic 98% B for 2.7 min, and then immediately stepped back down to 10% for 1.5 min re-equilibration. Flow rate was set at 0.5 mL/min. Compound MBQ-167 and the internal standard IPI549 were detected on mass spectrometer by using multiple reaction monitoring transitions 339.5→ 180.2 m/z and 529.2 → 326.3 m/z respectively. Mass spectrometry parameters were optimized. Collision energy (CE) and declustering potential (DP) were manually tuned using the instrument software.

### Cell culture and reagents

All immortalized adherent cell lines were maintained at 37°C in a humidified incubator with 5% CO₂ and cultured in medium supplemented with 10% fetal bovine serum (FBS; Cat# S11550, Atlanta Biologicals), 100 μg/mL Normocin (Cat# ant-nr-1, InvivoGen), and 100 U/mL Penicillin–Streptomycin–Glutamine (Cat# 10378-016, Gibco). NSCLC cell lines (H358, H460, and H520), Beas-2B immortalized normal human bronchial epithelium cell line, and HNC cell lines (Detroit 562, FaDu, SCC90) were all obtained from the American Type Culture Collection (ATCC). NSCLC cell lines were cultured in RPMI 1640 medium (Cat# 11875093, Gibco), whereas HNC cell lines were cultured in Eagle’s Minimum Essential Medium (EMEM; Cat# 30-2003, ATCC). Beas-2B cells were cultured in Airway Epithelial Cell Basal Medium (PCS-300-030, ATCC) and Bronchial Epithelial Cell Growth Kit (PCS-300-040, ATCC). Glioblastoma (GBM) cell lines (DBTRG, 8MG, and GBM38), which have been used and described in our previous studies (16,22), and normal human astrocytes, provided by Dr. Sofia Merajver at the University of Michigan (29), were all cultured in Dulbecco’s Modified Eagle Medium (DMEM; Cat# 11965-092, Gibco). Cell lines were authenticated by the originating repositories before distribution and were used immediately upon receipt. Cell lines maintained in continuous culture for more than one year were re-authenticated by short tandem repeat (STR) profiling. Mycoplasma testing was performed monthly using the MycoAlert™ Mycoplasma Detection Kit (Cat# LT07-418, Lonza).

### Compounds

MBQ-167 used for both *in vitro* cell-based assays and *in vivo* xenograft studies was provided by MBQ Pharma (30-32). The CHK1 inhibitor Rabusertib was purchased from Cayman Chemical (Cat# 20351). The CHK2 inhibitor PV1019 (Cat# HY-125203) and the NUAK1 inhibitor HTH-01-015 (Cat# HY-12334) were purchased from MedChemExpress.

### Plasmids and transfection

pcDNA3.1(+)-3xFlag-tagged Abi-1 wild type and site-directed mutants (Abi-1-S323D and Abi-1-S323A) were described as previously (22). pRK5-Myc-tagged Rac1 wild type (Cat# 12985), Q61L (Cat# 12983), and T17N (Cat# 12984) plasmids were obtained from Addgene. Plasmid DNA was transfected using Lipofectamine 2000 (Cat# 52887, Invitrogen) according to the manufacturer’s protocol.

### Irradiation

*In vitro* and animal irradiation experiments were conducted using a Philips RT250 irradiator (Kimtron Medical) at the Experimental Irradiation Core Facility of the University of Michigan Rogel Cancer Center (RRID: SCR_025766). Radiation output was verified using an ionization chamber coupled to an electrometer, with calibration traceable to standards established by the National Institute of Standards and Technology (NIST). For localized irradiation of flank tumors, mice were maintained under isoflurane anesthesia and arranged two at a time for treatment. The tumor-bearing area of each mouse was positioned at the center of a 2.4-cm aperture in the secondary collimator, while the remainder of the animal was shielded to minimize radiation exposure outside the treatment field.

### Immunofluorescence

Cells were seeded onto coverslips, treated under the indicated conditions, and fixed at the indicated time points with 4% paraformaldehyde. For γ-H2AX staining, cells were incubated with mouse anti-phospho-Histone H2AX antibody (1:1,000; Cat# 05-636, Millipore), followed by Alexa Fluor 594-conjugated goat anti-mouse IgG secondary antibody (1:2,000; Cat# A-11005, Invitrogen). DNA was counterstained with DAPI. γ-H2AX foci were quantified in at least 50 cells per condition, and cells containing ≥10 γ-H2AX foci were scored as positive. For 53BP1 and RAD51 staining, cells were incubated with rabbit anti-53BP1 antibody (1:500; Cat# 4937S, Cell Signaling Technology) and mouse anti-RAD51 antibody (1:100; Cat# GTX70230, GeneTex), followed by Alexa Fluor 488-conjugated goat anti-rabbit IgG (1:1,000; Cat# ab150077, Abcam) and Alexa Fluor 594-conjugated goat anti-mouse IgG (1:1,000; Cat# A-11005, Invitrogen). Nuclei were counterstained with DAPI. The foci threshold, which is used to define a positive cell, was 10 for 53BP1 and 5 for Rad51. Images were acquired using an ECLIPSE Ti2 inverted microscope (Nikon).

### Celltiter-Glo cell viability assay

Cells were seeded into 96-well plates at approximately 2,000 cells per well. After 24 hours, cells were treated with escalating concentrations of compounds (Rabusertib, PV1019, or HTH-01-015) and irradiated 2–4 hours after drug treatment. Five days later, cell viability was assessed using the CellTiter-Glo® 2D Cell Viability Assay (Cat# G9242, Promega) according to the manufacturer’s instructions.

### Neutral comet assay

H460 and Detroit 562 cells were plated and treated with indicated conditions at different time points. Single-cell gel electrophoretic comet assays were performed under Neutral conditions as described previously (16,33). Briefly, cells were combined with 1% LM Agarose (Cat# IB70051, IBI SCIENTIFIC) at 40 °C at a ratio of 1:10 (vol/vol) and immediately pipetted onto slides. For cellular lysis, the slides were immersed in lysis solution (Cat# 4250-050-01, R&D SYSTEMS) overnight at 4 °C in the dark, followed by washing in Neutral Electrophoresis Buffer (100 mM Tris, 300 mM sodium acetate, pH 9.0, adjusted with glacial acetic acid) for 30 min. Slides were then immersed in Neutral Electrophoresis Buffer to be subjected to electrophoresis at 21 V for 30 min and stained in SYBR™ Gold Nucleic Acid Gel Stain (Cat# S11494, Invitrogen) for 20 min. All images were taken with a fluorescence microscope and analyzed by Comet Assay IV software (Perceptive Instruments). For quantification, the tail moment, a measure of both amount and distribution of DNA in the tail, was used as an indicator of DNA damage. Comet-positive cells (50-100 cells) were scored in random fields for each experimental condition. Images were acquired using an ECLIPSE Ti2 inverted microscope (Nikon).

### Clonogenic survival assay

Cells were treated with or without the indicated compounds and seeded to 6-well plates at clonal density. After allowing cells to attach, they were irradiated with the indicated doses. Colonies were allowed to grow for 10–14 days (21 days for SCC90 cells), then fixed, stained, and counted. Colonies containing more than 50 cells were scored as surviving colonies. The radiation enhancement ratio (RER) was calculated as the ratio of the mean inactivation dose (Dmid) for the control group to that for the drug-treated group and was defined as the area under the fitted linear-quadratic clonogenic survival curve (16,34).

### Immunoblotting assay

Cells or xenograft tumor tissues were ground and lysed using RIPA lysis buffer (Cat# 89900, Thermo Scientific) supplemented with PhosSTOP phosphatase inhibitor (Cat# 04906845001, Roche) and complete protease inhibitor tablets (Cat# 1187358001, Roche). Proteins were detected with primary antibodies (1:1000 dilution) for γ-H2AX (Cat# 05-636, Millipore), Myc-tag (Cat# SAB4700447, Sigma-Aldrich), Flag-tag (Cat# F1804, Sigma-Aldrich), β-Actin (Cat# sc-47778, Santa Cruz Biotechnology), Abi-1 (Cat# sc-398554, Santa CruzBiotechnology), Rac1 (Cat# ARC03, Cytoskeleton Inc.), p-CHK1 (Ser296) (Cat# 2349, Cell signaling), CHK1 (Cat# 2360, Cell signaling), p-CHK2(Ser516) (Cat# 2669, Cell signaling), CHK2 (Cat# 3440, Cell signaling), p-MYPT1(Ser668) (Cat# 3048, Cell signaling), and MYPT1 (Cat# 2634, Cell signaling) and second antibodies (1:3000 dilution) of rabbit and mouse IgG. Rabbit polyclonal p-Abi-1 (Ser323) antibody was raised by Cell Signaling Technology as described previously (22).

### GTP-Rac1 activity Assay

GTP-bound Rac1 activity was measured as previously described (22), according to the manufacturer’s protocol (Cytoskeleton Inc., Cat# BK035). Briefly, approximately 500-800 μg of total protein from cell lysates or tumor lysates was incubated with 10–20 μl of PAK-PBD agarose beads at 4°C for 1 hour to selectively pull down active GTP-Rac1. Following incubation, beads were washed, and bound proteins were eluted in 20 μl of 2× Laemmli sample buffer (Bio-Rad, Cat# 161-0737). Eluted samples were subjected to immunoblotting using an anti-Rac1 antibody (Cytoskeleton Inc., Cat# ARC03).

### Immunohistochemical staining

Mouse tumor tissues were harvested, fixed in 10% neutral-buffered formalin, and embedded in paraffin. Protein expression in FFPE mouse tumor tissues and patient TMAs was evaluated by immunohistochemistry (IHC). IHC was performed using the ABC Vectastain Kit (Vector Laboratories) according to the manufacturer’s instructions. Briefly, paraffin-embedded tissue sections were deparaffinized, rehydrated, subjected to antigen retrieval, and blocked before incubation with a primary antibody against γ-H2AX (1:1,000 dilution; Millipore, Cat# 05-636), or p-Abi-1 (1:200 dilution; Cell Signaling(22)) at 4°C overnight. Following incubation with the appropriate secondary antibody for 30 minutes, sections were developed using 3,3′-diaminobenzidine (DAB) and counterstained with hematoxylin. IHC quantification of p-Abi-1 was performed as described previously (22). IHC staining for γ-H2AX was quantified by analyzing the percentage of positive cells in five representative fields from each tumor. Briefly, the percentage of γ-H2AX-positive cells was calculated as the number of positively stained cells divided by the total number of cells. The percentages from the five fields were averaged to generate a single value for each tumor. Each data point represents one individual tumor. Images were acquired using an ECLIPSE Ti2 inverted microscope (Nikon).

### Statistical Analysis

Statistical analyses of γ-H2AX foci formation, clonogenic survival, mean inactivation dose (Dmid), and comet assay data were performed using two-tailed Student’s *t*-tests in GraphPad Prism version 11. For xenograft studies, tumor volumes were normalized to 100% at the initiation of treatment for each experimental group. Tumor volume over time was compared among treatment groups in both H460 and Detroit 562 xenograft models using two-way ANOVA. Time to tumor doubling or sextupling was defined as the earliest time point at which tumor volume reached at least two-fold (Detroit 562) or six-fold (H460) of the baseline volume at the start of treatment. These endpoints were analyzed using Kaplan–Meier survival curves and compared between groups using the log-rank (Mantel–Cox) test. The overall survival (OS) or disease specific survival (DSS) in GBM and LSCC cohorts were determined by Kaplan-Meier analysis stratifying by p-Abi-1 status. The chi-square test or Fisher’s exact test was employed to evaluate the relationship between p-Abi-1-S323 and clinicopathological variables. The analysis was done with SPSS software (SPSS v. 22.0). A p value < 0.05 was considered statistically significant.

### Data Availability

The data generated in this study are available upon request from the corresponding author.

## Results

### Abi-1-S323 dephosphorylation predicts poor patient outcomes following RT

Since our previous studies established that Rac1-Abi-1 signaling regulates genotoxic treatment response in preclinical models of GBM, we next sought to determine its relevance in human patients and across diseases. We established a tissue microarray (TMA) from primary glioma specimens, including 33 GBM patients who received RT as part of their clinical care (23), with specimen collected before RT. We assessed tumor phosphorylated Abi-1-S323 (p-Abi-1-S323) by immunohistochemistry, classifying tumors as p-Abi-1-positive or -negative as previously described (**Fig. 1A**) (22) and correlated this staining with clinical outcomes. Patients with p-Abi-1-positive tumors had significantly longer overall survival (OS) than those with p-Abi-1-negative tumors (22.8 ± 1.3 vs. 8.9 ± 1.9 months; *p* = 0.002; **Fig. 1B**). We found no significant association between p-Abi-1-S323 status and other patient characteristics (**Table 1**), including MGMT status. These data support the concept that dephosphorylated Abi-1-S323 promotes DNA repair and RT resistance in glioma patients and prompting us to evaluate its role in RT responses across human cancers.

**Table 1.** Association of p-Abi-1-S323 expression with patient characteristics in GBM.

|  | All case | p-Abi-1 |  | p |
| --- | --- | --- | --- | --- |
|  |  | Positive | Negative |  |
| **Age** |  |  |  |  |
| ≤ 58 | 17 | 13 | 4 | **0.218** |
| > 58 | 16 | 9 | 7 |  |
| **Gender\*** |  |  |  |  |
| Male | 10 | 7 | 3 | **1.000** |
| Female | 22 | 15 | 7 |  |
| **MGMT** |  |  |  |  |
| Methylated | 11 | 8 | 3 | **0.709** |
| Unmethylated | 22 | 14 | 8 |  |
| **Extent of resection** |  |  |  |  |
| STR (Subtotal Resection) | 8 | 5 | 3 | **1.000** |
| GTR/NTR<br>(Gross/Near Total Resection) | 25 | 17 | 8 |  |
| **Performance status** |  |  |  |  |
| 0 | 3 | 2 | 1 | **0.548** |
| 1 | 17 | 12 | 5 |  |
| 2 | 12 | 8 | 4 |  |
| 3 | 1 | 0 | 1 |  |

Note: \* indicates missing patient information for the variable, resulting in a total number of patients of less than 33.

Since Abi-1 phosphorylation is a novel post-translational mark that cannot be assessed in large tumor sequencing datasets, we focused on Rac1 to identify other cancer types with potentially elevated Rac1-Abi-1 signaling. We analyzed RNA-sequencing data from cBioPortal (September 2024) and found that *RAC1* was most frequently amplified in NSCLC, bladder cancer, and HNC (**Fig. 1C**). Because RT is a standard treatment for patients with both NSCLC and HNC, we selected these cancer types for further investigation. Analysis of *RAC1* mRNA expression and clinical outcomes in The Cancer Genome Atlas (TCGA) further showed that lower *RAC1* expression was associated with longer OS in both NSCLC and HNC (Supplementary **Fig. S1A**-**B**), supporting a potential role for Rac1 signaling in these cancers. We therefore examined p-Abi-1-S323 in an independent HNC TMA cohort (24,25) (**Fig. 1D**). Among 72 patients who received RT as part of their clinical care, those with pre-treatment p-Abi-1-positive tumors had significantly longer OS (p = 0.02; **Fig. 1E**) and disease-specific survival (DSS; p = 0.03; **Fig. 1F**) compared to those with p-Abi-1-negative tumors. There was no significant association between p-Abi-1-S323 status and other patient characteristics (**Table 2**), suggesting that patients whose tumors harbor Abi-1 in an inactive phosphorylated state may have improved responses to RT. Together, these findings suggest that p-Abi-1-S323 status may predict patient survival and support a clinically relevant role for Rac1-Abi-1 signaling in RT response in human cancers, particularly in NSCLC and HNC.

**Table 2.** Association of p-Abi-1-S323 expression with patient characteristics in HNC.

|  | All case | p-Abi-1 |  | p |
| --- | --- | --- | --- | --- |
|  |  | Positive | Negative |  |
| **Age** |  |  |  |  |
| ≤ 61 | 41 | 27 | 14 | **0.342** |
| >61 | 31 | 17 | 14 |  |
| **Gender** |  |  |  |  |
| Male | 56 | 36 | 20 | **0.301** |
| Female | 16 | 8 | 8 |  |
| **Tobacco\*** |  |  |  |  |
| Never | 4 | 2 | 2 | **0.614** |
| Former | 29 | 16 | 13 |  |
| Current | 38 | 25 | 13 |  |
| **ACE Comorbidity Index** |  |  |  |  |
| None/Mild | 45 | 27 | 18 | **0.803** |
| Mod/Severe | 27 | 17 | 10 |  |
| **Site Supraglottic\*** |  |  |  |  |
| Supraglottic | 45 | 28 | 17 | **0.984** |
| Not | 16 | 10 | 6 |  |
| **Overall Stage** |  |  |  |  |
| 2 | 1 | 1 | 0 | **0.604** |
| 3 | 20 | 11 | 9 |  |
| 4 | 51 | 32 | 19 |  |
| **Node Invasion\*** |  |  |  |  |
| 0 (no) | 16 | 10 | 6 | **0.984** |
| 1 (yes) | 45 | 28 | 17 |  |

| Differentiation* |  |  |  |  |
| --- | --- | --- | --- | --- |
| Moderate | 34 | 23 | 11 | **0.407** |
| Moderate to Poor | 1 | 0 | 1 |  |
| Poor | 17 | 8 | 9 |  |
| Well | 5 | 4 | 1 |  |
| Well to Moderate | 1 | 1 | 0 |  |

Note: \* indicates missing patient information for the variable, resulting in a total number of patients of less than 72.

### Rac1-Abi-1 signaling is elevated in NSCLC and HNC and correlates with radioresistance

To further interrogate the causal links between Rac1-Abi-1 and RT resistance in NSCLC and HNC, we measured Rac1 activity using a GTP-Rac1 pull-down assay followed by immunoblotting as described previously (22), and evaluated radiosensitivity in models of both cancer types. Among the NSCLC cell lines examined, H520 cells exhibited the lowest Rac1 activity and highest Abi-1-S323 phosphorylation (**Fig. 2A-C**) and were the most radiosensitive, as indicated by reduced clonogenic survival (**Fig. 2D**) and the lowest mean inactivation dose (Dmid) (Supplementary **Fig. S2A**). In contrast, H460 cells exhibited the highest Rac1 activity (H460 vs. H520 densitometric values: 0.79 ± 0.21 vs. 0.02 ± 0.02; p = 0.003; **Fig. 2B**), lowest Abi-1-S323 phosphorylation (0.07 ± 0.07 vs. 0.88 ± 0.10; p < 0.0001; **Fig. 2C**), and greatest radioresistance (**Fig. 2D**; Dmid: 3.03 ± 0.30 vs. 1.52 ± 0.29; p = 0.001; Supplementary **Fig. S2A**). H358 cells exhibited intermediate Rac1 activity and p-Abi-1-S323 levels and correspondingly intermediate radiosensitivity. Because TP53 status is an important determinant of cellular responses to DNA damage and therapeutic efficacy in NSCLC (35), we considered whether it might account for these differences. Notably, H460 cells are TP53 wild-type, H520 cells harbor mutant TP53, and H358 cells are TP53-null (36). Rac1-Abi-1 signaling activity consistently correlated with RT sensitivity across these distinct TP53 backgrounds, suggesting that this association is not simply explained by TP53 status. These findings support Rac1-Abi-1 signaling as an independent determinant of RT sensitivity in NSCLC.

**Figure 2.**
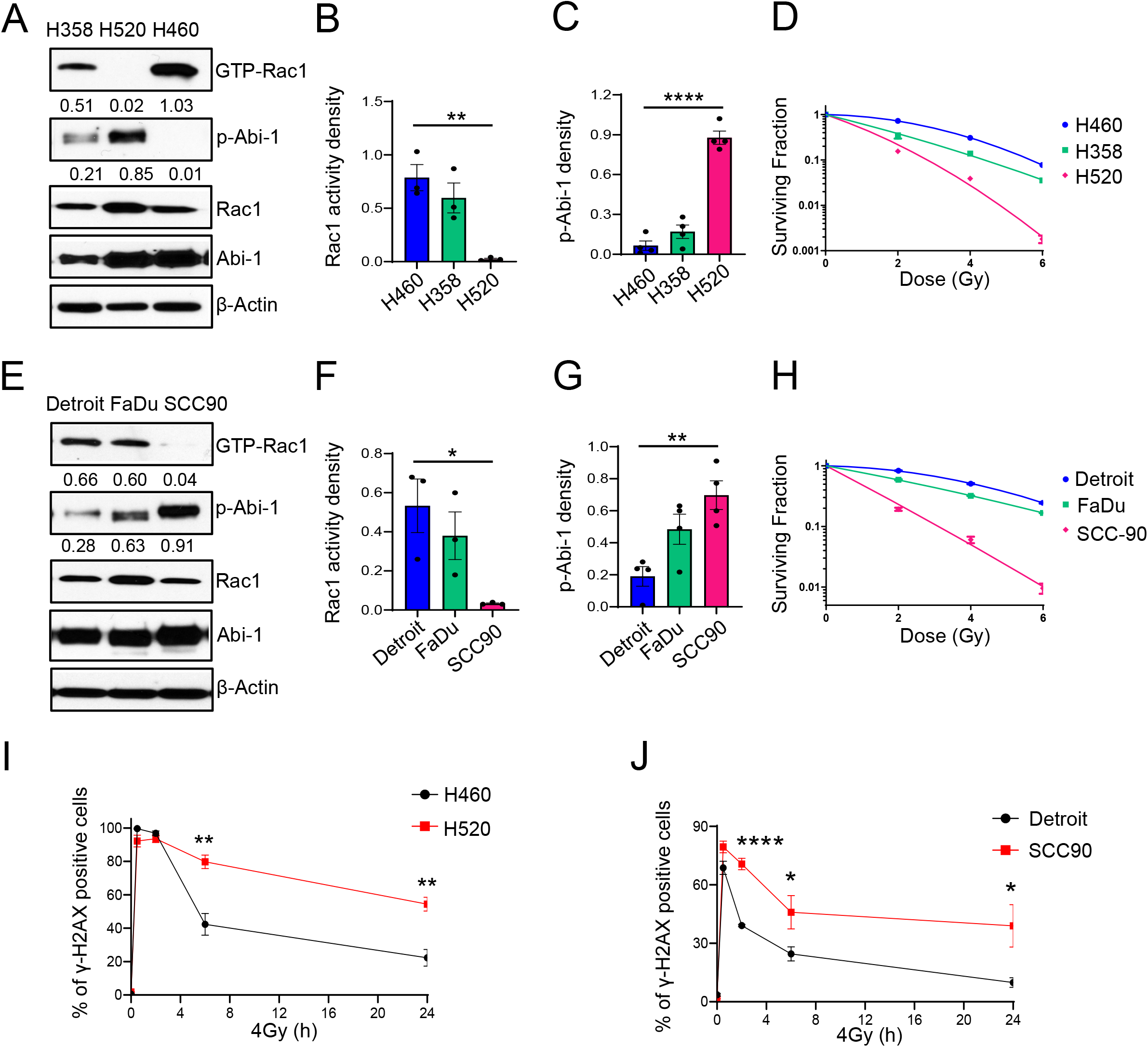
High Rac1-Abi-1 signaling activity correlates with radioresistance in NSCLC and HNC. **(A-C)** GTP-bound Rac1 activity and p-Abi-1-S323 levels in the NSCLC cell lines H358, H520, and H460 were determined by GTP-Rac1 pull-down assay and immunoblotting, respectively. Densitometric analysis was performed using ImageJ, and values are shown below the corresponding bands. Representative immunoblots are shown in **(A)**, with quantification of GTP-bound Rac1 activity **(B),** and p-Abi-1-S323 levels **(C)** from 3-4 independent biological replicates. **, p < 0.01; ****, p < 0.0001. **(D)** Three NSCLC cells were irradiated with escalating doses of radiation, and colonies were stained and quantified 10-14 days after RT. Representative clonogenic survival curves from 3-5 independent biological replicates are shown. **(E-G)** GTP-bound Rac1 activity and p-Abi-1-S323 levels in the HNC cell lines Detroit 562 (Detroit), FaDu, and SCC90 were determined as described above. Representative immunoblots are shown in **(E)**, with quantification from 3-4 independent biological replicates shown in (**F-G**). * p < 0.05; **, p < 0.01. **(H)** Three HNC cells were irradiated with escalating doses of radiation, and colonies were stained and quantified 10-14 days after RT. Representative clonogenic survival curves from 3-5 independent biological replicates are shown. **(I-J)** Cells of the indicated lines were irradiated with 4 Gy and fixed 0.5, 2, 6, 24 hours later for immunofluorescence staining of γ-H2AX. Quantification of γ-H2AX foci from 3-4 independent biological replicates is shown. Scale bar, 5 μm. *, p < 0.05; **, p < 0.01; ****, p < 0.0001. For **(B)**, **(C)**, **(F)**, **(G)**, **(I)**, and **(J)**, data are presented as mean ± SEM and were analyzed using a two-tailed t test.

A similar pattern was observed in HNC cell lines. SCC90 cells displayed the lowest Rac1 activity (**Fig. 2E** and **2F**), the highest Abi-1 phosphorylation (**Fig. 2E** and **2G**), and the greatest radiosensitivity (**Fig. 2H**; Supplementary **Fig. S2B**). In contrast, Detroit 562 cells exhibited the highest Rac1 activity (Detroit 562 vs SCC90: 0.53 ± 0.24 vs 0.03 ± 0.006; p = 0.02; **Fig. 2F**), the lowest Abi-1 phosphorylation (0.12 ± 0.0.24 vs 0.77 ± 0.18; p = 0.003; **Fig. 2G**), and were the most resistant to RT (**Fig. 2H**; Dmid: 4.02 ± 0.31 vs 1.53 ± 0.17; p < 0.0001; Supplementary **Fig. S2B**). FaDu cells exhibited intermediate Rac1 activity and p-Abi-1-S323 levels and showed a corresponding intermediate response to RT. Human papillomavirus (HPV) status is a well-established determinant of RT sensitivity in HNC(37). Although HPV-positive SCC90 cells exhibited lower Rac1-Abi-1 signaling activity and greater RT sensitivity, the HPV-negative Detroit 562 and FaDu cells displayed distinct Rac1-Abi-1 signaling activities. These findings suggest that Rac1-Abi-1 signaling may represent an additional determinant of RT response in HNC.

Based on these findings, we selected H460/H520 and Detroit 562/SCC90 as paired NSCLC and HNC models, respectively, for subsequent studies. To determine whether RT sensitivity correlated with DNA repair, we quantified residual γ-H2AX foci, a surrogate marker of DSBs (16,22), following RT. Consistent with the clonogenic survival results, radioresistant H460 and Detroit 562 cells exhibited fewer residual γ-H2AX foci, indicating more efficient DSB repair, whereas radiosensitive H520 and SCC90 cells retained significantly more γ-H2AX foci (**Fig. 2I** and **2J**; Supplementary **Fig. S2C** and **S2D**). Thus, elevated Rac1 activity and reduced Abi-1 phosphorylation were associated with enhanced DSB repair and RT resistance in both NSCLC and HNC.

### Genetic modulation of Rac1 activity and Abi-1 phosphorylation regulates radioresistance in NSCLC and HNC

To determine whether Rac1 activation and Abi-1 dephosphorylation causally regulate RT response in NSCLC and HNC, we genetically manipulated Rac1 activity and Abi-1-S323 phosphorylation. In radioresistant H460 and Detroit 562 cells, which exhibit high Rac1 activity and low Abi-1 phosphorylation, expression of dominant-negative Rac1-T17N or phospho-mimetic Abi-1-S323D significantly increased RT sensitivity, as indicated by reduced clonogenic survival and increased residual γ-H2AX foci following RT (**Fig. 3A-F**; Supplementary **Fig. S3A** and **S3B**). Conversely, expression of constitutively active Rac1-Q61L or dephospho-mimetic Abi-1-S323A in radiosensitive H520 and SCC90 cells, which exhibit low Rac1 activity and high Abi-1 phosphorylation, increased clonogenic survival and reduced γ-H2AX foci following RT (**Fig. 3G-L**; Supplementary **Fig. S3C** and **S3D**). These findings establish Rac1-Abi-1 signaling as a causal determinant of RT response in NSCLC and HNC.

**Figure 3.**
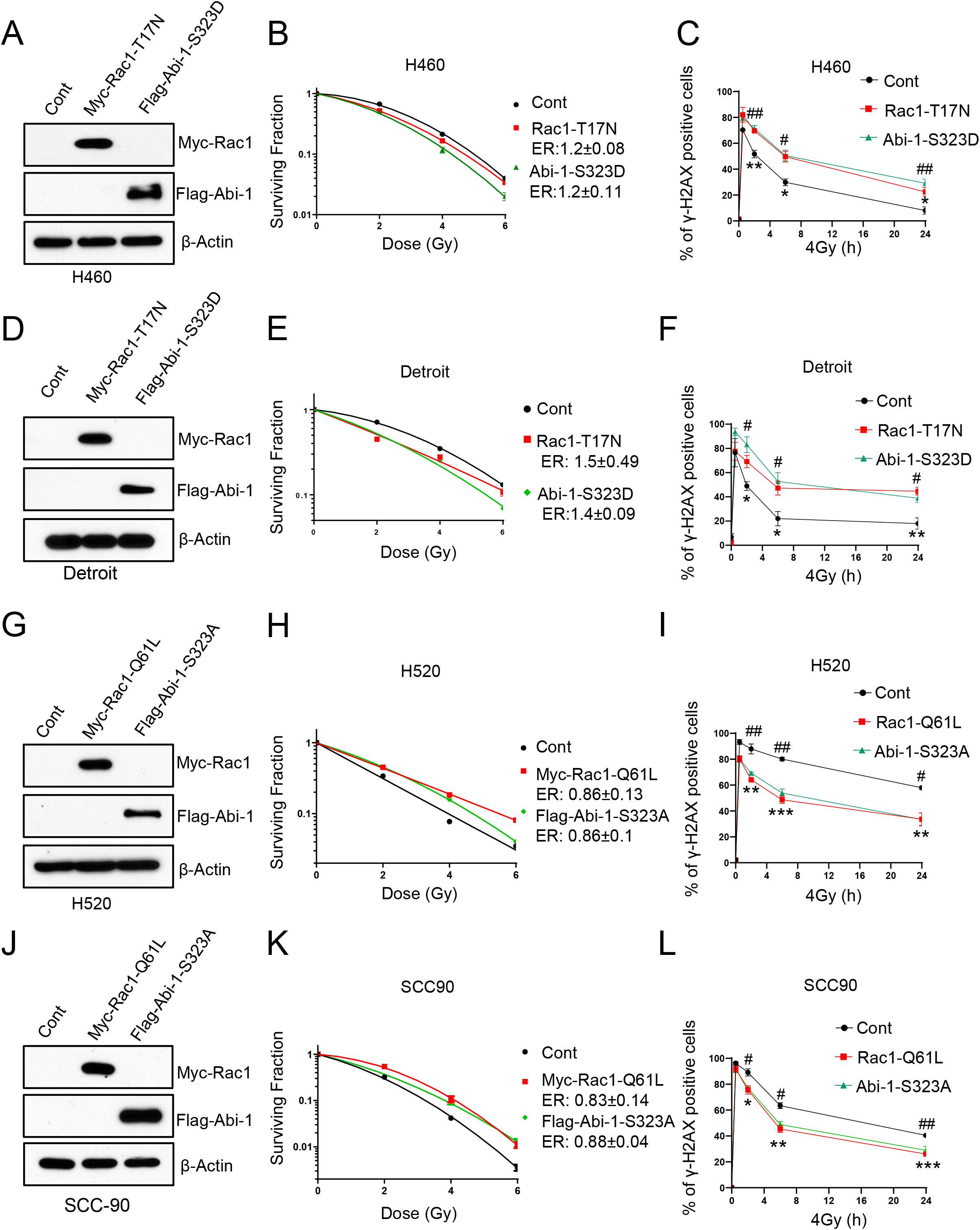
Genetic modulation of Rac1 activity and Abi-1 phosphorylation regulates DNA repair and radioresistance in NSCLC and HNC. **(A-F)** H460 and Detroit 562 (Detroit) cells were transiently transfected with myc-tagged Rac1-T17N or Flag-tagged Abi-1-S323D, followed by IB to confirm protein expression **(A, D)**. Transfected cells were then replated for clonogenic survival assays following irradiation **(B, E),** or fixed 0.5, 2, 6, 24 hours after irradiation (4 Gy) for γ-H2AX IF staining and quantification **(C, F)**. ER in **(B)** and **(E)** indicates the Enhancement Ratio, which is the ratio of Dmid control and Dmid treatment. ER > 1 indicates radiosensitization while ER < 1 indicates radioprotection. ER (mean ± SD) from the three biologic replicates is shown. For **(C)** and **(F)**, ‘ * ’ indicates comparisons between the control and Rac1-T17N groups at the corresponding time point, whereas ‘ # ’ indicates comparisons between the control and Abi-1-S323D groups. **(G-L)** H520 and SCC90 cells were transiently transfected with myc-tagged Rac1-Q61L or Flag-tagged Abi-1-S323A, followed by IB **(G, J)**, clonogenic survival assays **(H, K)**, or γ-H2AX IF staining and quantification **(I, L)** as above. Enhancement ratios (ERs) were calculated from 3-4 independent biological replicates for **(H)** and **(K)**. ‘ * ’ indicates comparisons between the control and Rac1-Q61L groups at the corresponding time point, whereas ‘ # ’ indicates comparisons between the control and Abi-1-S323A groups for **(I)** and **(L)**. For **(C)**, **(F)**, **(I)**, **and (L)**, * or #, p < 0.05; ** or ##, p < 0.01; *** or ###, p < 0.001. Data are presented as mean ± SEM (two-tailed t test).

### Pharmacological inhibition of Rac1-Abi-1 signaling blocks DSB repair and reverses radioresistance in NSCLC and HNC cells

We next asked whether pharmacological inhibition of the Rac1-Abi-1 pathway could similarly enhance RT response. We employed MBQ-167, a dual Rac1/CDC42 inhibitor with preclinical antitumor (31,38) and radiosensitizing activity (22) that is currently undergoing Phase I clinical evaluation in patients with advanced breast cancer (NCT06075810). Recent studies suggest that Rac1 inhibition contributes predominantly to its antitumor activity (32). In Rac1-Abi-1-active, radioresistant H460 and Detroit 562 cells, RT induced Rac1 activation and reduced Abi-1 phosphorylation, whereas MBQ-167 blocked these effects (**Fig. 4A** and **4B**). Consistently, MBQ-167 significantly enhanced RT sensitivity, as demonstrated by reduced clonogenic survival (**Fig. 4C** and **4D**), without radiosensitizing normal cells (Supplementary **Fig. S4A** and **S4B**), suggesting that tumor cells may exhibit a greater dependency on Rac1-Abi-1 signaling. MBQ-167 also significantly increased residual γ-H2AX foci (**Fig. 4E** and **4F**; Supplementary **Fig. S4C** and **S4D**) and delayed DSB resolution, as reflected by increased tail moments in the neutral comet assay (**Fig. 4G** and **4H**) in both models. Together, these findings demonstrate that pharmacological inhibition of Rac1-Abi-1 signaling impairs DSB repair and selectively enhances RT response in Rac1-Abi-1-active cancer cells.

**Figure 4.**
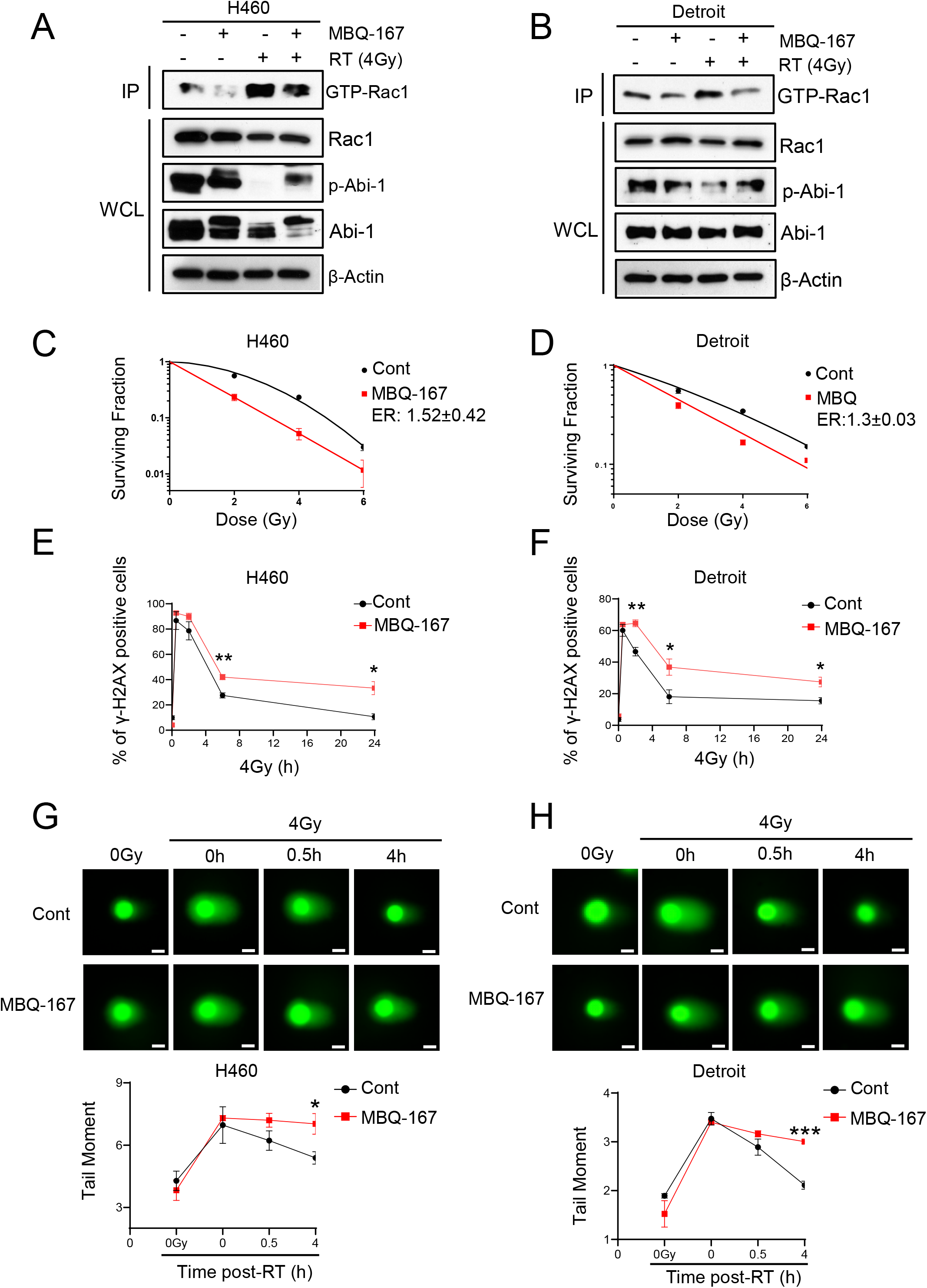
Pharmacological inhibition of Rac1-Abi-1 signaling enhances radiosensitivity by suppressing DNA DSB repair. **(A-B)** H460 and Detroit 562 (Detroit) cells were treated with MBQ-167 (100 nM) overnight. Cells were harvested to determine GTP-bound Rac1 activity and p-Abi-1-S323 levels. Representative data from 3-5 biological repeats is shown. IP: immunoprecipitation; WCL: whole cell lysates. **(C-D)** H460 and Detroit cells were treated with MBQ-167 (100 nM) overnight and then replated for clonogenic survival assays following RT. Representative clonogenic survival curves from 3 independent biological replicates are shown. ER (mean ± SD) from the three biologic replicates is shown. **(E-F)** H460 and Detroit cells were treated with MBQ-167 overnight. Cells were replated and fixed for γ-H2AX IF staining 0.5, 2, 6, 24 hours after RT (4Gy), and γ-H2AX foci were quantified. **(G-H)** H460 and Detroit cells were treated with MBQ-167 overnight and harvested at the indicated time points following irradiation for neutral comet assays. Tail moment (bottom panels), which reflects both the amount of DNA in the comet tail and the distance of DNA migration, was used as an indicator of DNA damage. For the 0-hour time point, cells were irradiated (4 Gy) and harvested on ice to prevent DNA repair, while 0.5-hour and 4-hour time point were processed at room temperature. Scale bar: 20 μm. For **(E)**, **(F)**, **(G)**, and **(H)**, *, p < 0.05; **, p < 0.01, ***, p < 0.001, comparing control and MBQ-167 groups at the corresponding time point. Data are presented as mean ± SEM from 3-4 independent biological replicates and were analyzed using a two-tailed *t* test.

### Pharmacological inhibition of Rac1-Abi-1 signaling suppresses tumor growth and prolongs mouse survival in NSCLC and HNC xenograft models

Given the robust radiosensitizing effects of MBQ-167 *in vitro*, we next evaluated its activity in RT-resistant xenograft models of H460 NSCLC and Detroit 562 HNC. Tumor-bearing mice were assigned to four treatment groups as described previously (22): vehicle control, MBQ-167 alone, RT alone, or the combination of MBQ-167 and RT. To assess drug exposure (pharmacokinetics, PK), target engagement, signaling changes, and DNA repair (pharmacodynamics, PD), mice received an abbreviated treatment regimen consisting of two daily fractions of RT (2 Gy/fraction) and three daily doses of MBQ-167 (75 mg/kg/day), and tumors were harvested 4 hours after the final RT fraction (Supplementary **Fig. S5A** and **S5D**, upper timelines).

PK analysis demonstrated accumulation of MBQ-167 in both plasma (**Fig. 5A** and **5G**) and tumors (**Fig. 5B** and **5H**) in mice treated with MBQ-167 alone or in combination with RT in both models. PD analyses showed that RT activated Rac1-Abi-1 signaling, as evidenced by increased GTP-bound Rac1 and decreased Abi-1 S323 phosphorylation, whereas MBQ-167 blocked these RT-induced changes (**Fig. 5C** and **5I**). Consistent with Rac1-Abi-1 inhibition, MBQ-167 delayed repair of RT-induced DSBs, as demonstrated by increased γ-H2AX detected by immunoblotting (**Fig. 5C** and **5I**) and immunohistochemistry (**Fig. 5D** and **5J**; Supplementary **Fig. S5B** and **S5E**). These findings indicate that RT-induced Rac1 activation promotes Abi-1-S323 dephosphorylation and facilitates DSB repair *in vivo*.

**Figure 5.**
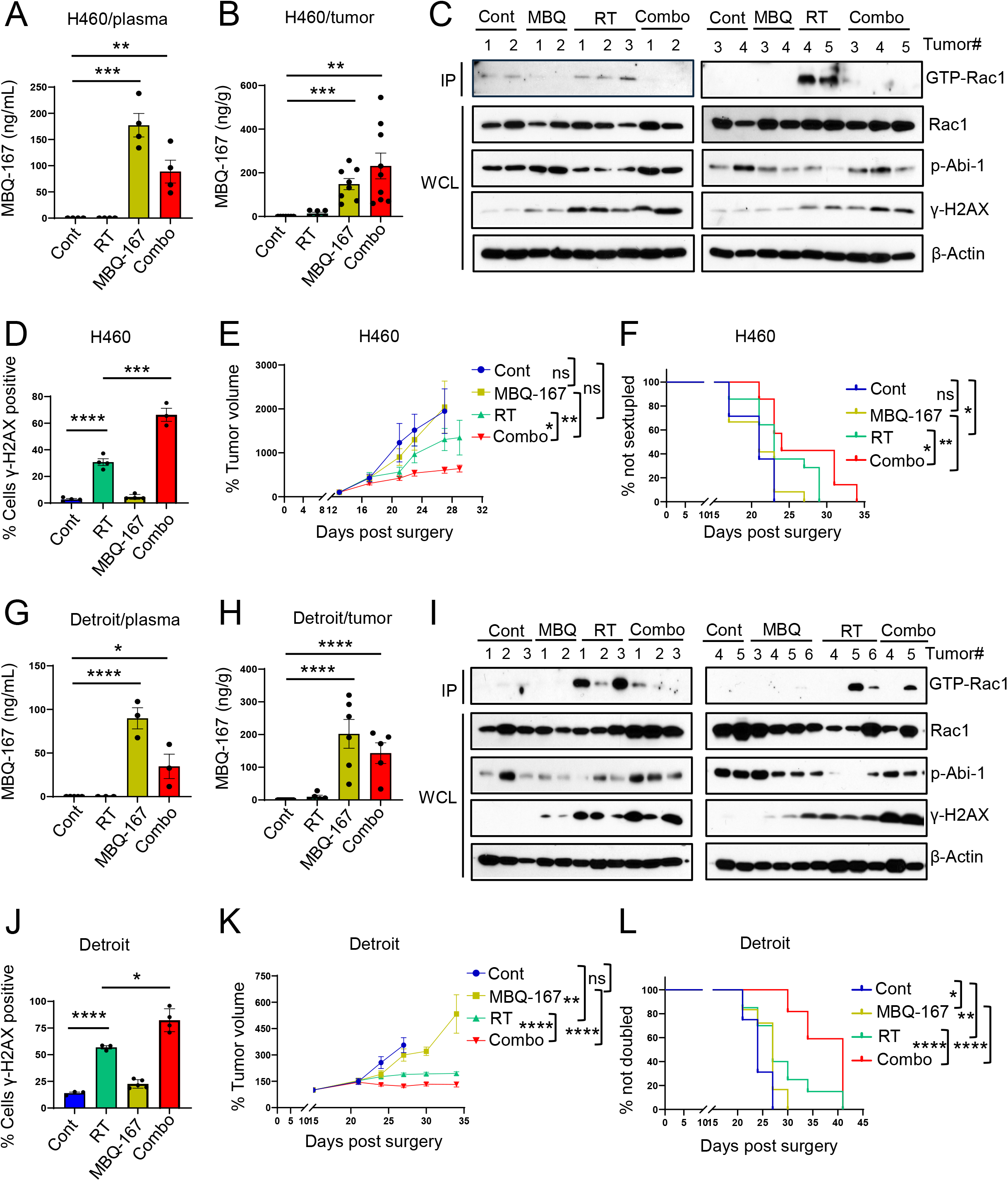
Pharmacological inhibition of Rac1-Abi-1 signaling enhances radiosensitivity in xenograft tumor models. **(A-D)** H460 flank xenograft tumors were established in mice. A subset of animals (3-5 mice/group; Supplementary **Fig. S5A**, above timeline) received three doses of the Rac1 inhibitor MBQ-167 and two fractions of RT (2 Gy/fraction). Tumors were harvested at ∼4 hours after the second radiation dose for PK analysis to evaluate drug delivery **(A, B)**, IB to assess target protein expression **(C)**, and IHC staining for γ-H2AX **(D)**. Five tumors from each treatment arm were analyzed by IB in **(C)**, and γ-H2AX levels were quantified in tumors from each treatment arm in **(D)**. **, p < 0.01; ***, p < 0.001; ****, p < 0.0001; mean ± SEM (two tailed t test). **(E-F)** A separate cohort of H460 tumor-bearing mice (n = 6-7 mice/group) received 12 doses of MBQ-167 and 10 fractions of radiation (2 Gy/fraction) to evaluate therapeutic efficacy (Supplementary **Fig. S5A**, bottom timeline). Tumor volumes were normalized to the individual tumor volume on Day 1 and are presented as mean ± SEM **(E)**. Kaplan–Meier analysis of time to tumor sixfold volume increase is shown **(F)**. **(G-J)** Detroit 562 (Detroit) flank xenograft tumors were established in mice. A subset of animals (3-5 mice/group; Supplementary **Fig. S5D**, above timeline) received the same short-term treatment regimen as described for the H460 model, and tumors were harvested for PK analysis, IB, and γ-H2AX IHC staining. Five to six tumors from each treatment arm were analyzed by IB in **(I)**. *, p < 0.05; ****, p < 0.0001; mean ± SEM (two tailed t test). **(K-L)** A separate cohort of Detroit tumor-bearing mice (n = 8-11 mice/group; Supplementary **Fig. S5D**, bottom timeline) received eight doses of MBQ-167 and six fractions of radiation (2 Gy/fraction) to evaluate therapeutic efficacy. Tumor volumes were normalized to the individual tumor volume on Day 1 and are presented as mean ± SEM **(K)**. Kaplan–Meier analysis of time to tumor doubling is shown **(L)**. Two-way ANOVA was used for **(E)** and **(K),** and the Logrank (Mantel-Cox) test was used for **(F)** and **(L)**. *, p < 0.05; **, p < 0.01, ****, p < 0.0001, and ns, no significant.

We next evaluated whether these molecular effects translate to therapeutic benefit. H460 tumor-bearing mice received 10 fractions of RT and/or 12 doses of MBQ-167 (Supplementary **Fig. S5A**, lower timeline), whereas Detroit 562 tumor-bearing mice received 6 fractions of RT and/or 8 doses of MBQ-167 (Supplementary **Fig. S5D**, lower timeline). In both models, MBQ-167 or RT alone moderately delayed tumor growth and prolonged survival, whereas the combination produced significantly greater tumor growth inhibition and survival benefits than either monotherapy in H460 NSCLC (**Fig. 5E** and **5F**) and Detroit 562 HNC (**Fig. 5K** and **5L**) models. MBQ-167 was well tolerated, with minimal effects on body weight throughout treatment (Supplementary **Fig. S5C** and **S5F**). Collectively, these findings demonstrate that pharmacological Rac1 inhibition enhances the therapeutic efficacy of RT across RT-resistant NSCLC and HNC models.

### CHK1 mediates Abi-1-S323 phosphorylation across multiple cancer models

We previously demonstrated that GTP-bound Rac1 activates protein phosphatase 5 (PP5) to mediate the dephosphorylation of Abi-1 at Ser323 (22). However, the kinase responsible for Abi-1-S323 phosphorylation remains unknown. Identifying this upstream kinase would provide important mechanistic insight into the regulation of Rac1-Abi-1 signaling and help distinguish the biological functions of Abi-1 phosphorylation and dephosphorylation under different cellular contexts. To identify candidate kinases, we leveraged a recently published kinome-wide kinase–substrate prediction resource that systematically characterized the substrate specificities of more than 300 human serine/threonine kinases and computationally predicted kinase–phosphosite relationships across the human phosphoproteome (39). This analysis identified checkpoint kinase 1 (CHK1), checkpoint kinase 2 (CHK2), and NUAK family kinase 1 (NUAK1/ARK5) as the top three candidate kinases predicted to phosphorylate Abi-1 at S323 (Supplementary **Fig. S6A**).

To validate these candidate kinases, we selected the GBM cell line DBTRG in which we originally identified Rac1-Abi-1 signaling pathway (22), the NSCLC cell line H520, and the HNC cell line SCC90, which exhibit relatively high p-Abi-1-S323 levels (Supplementary **Fig. S6B**). We treated these cells with selective inhibitors targeting the three top candidates: CHK1 (Rabusertib(40,41)), CHK2 (PV1019(42,43)), and NUAK1 (HTH-01-015(44)). Cells were then treated with escalating concentrations of each inhibitor based on their respective growth inhibition 50% (GI50) values (Supplementary **Fig. S6C–K**), followed by immunoblotting to assess p-Abi-1-S323 and target engagement. CHK1 inhibition consistently reduced p-Abi-1-S323 levels in all three cell lines (**Fig. 6A**), whereas CHK2 or NUAK1 inhibition had no detectable effect on Abi-1 phosphorylation (**Fig. 6B** and **6C**). Because protein phosphorylation is highly dynamic, we next performed a time-course experiment using a fixed concentration of the CHK1 inhibitor that effectively suppressed CHK1 activity and p-Abi-1 S323. CHK1 inhibition consistently reduced p-Abi-1-S323 levels at or after 2 h in all three cell lines (**Fig. 6D**), further supporting CHK1 as the kinase regulating Abi-1-S323 phosphorylation and establishing a previously unrecognized CHK1-Abi-1 signaling axis.

**Figure 6.**
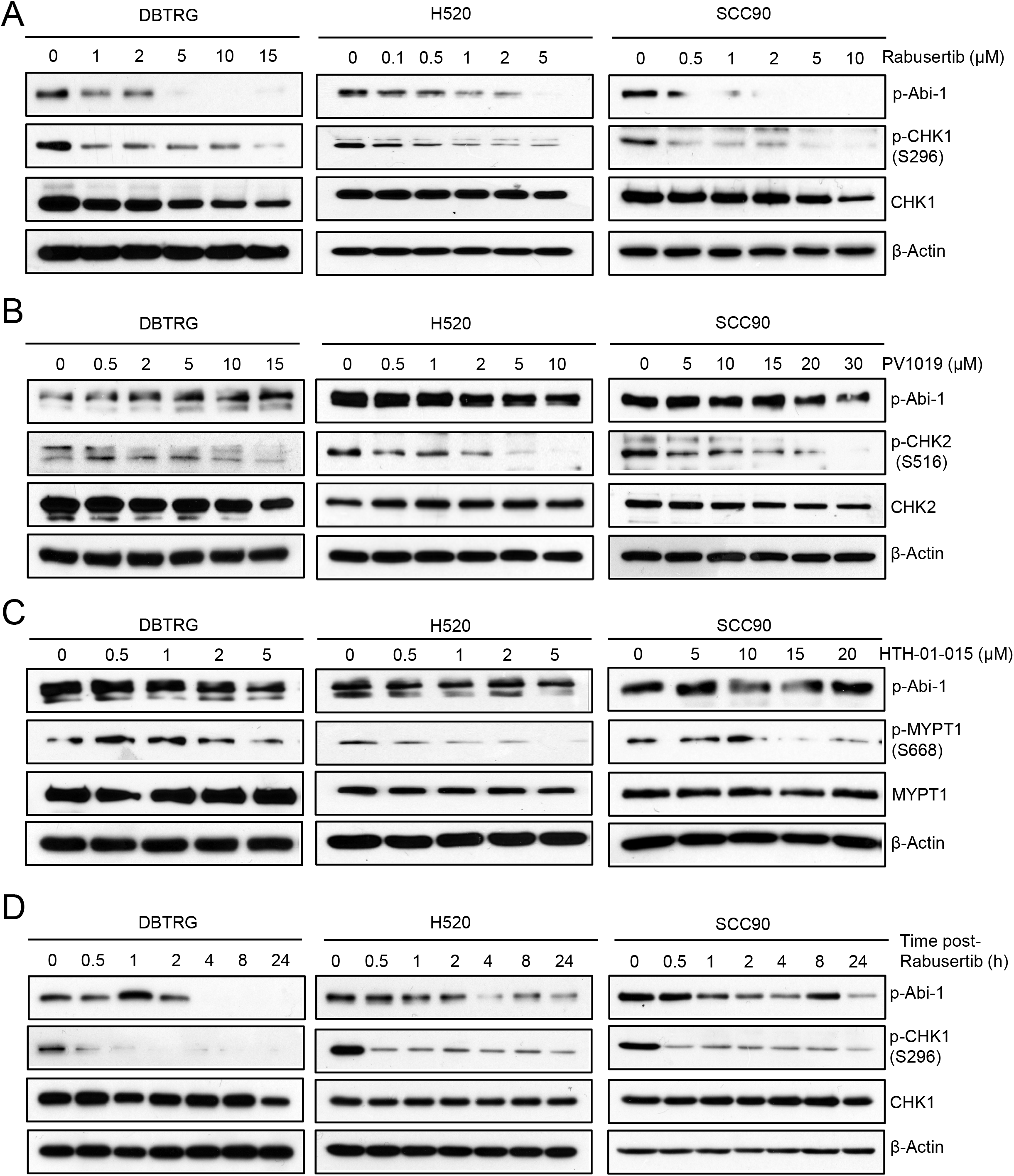
CHK1 kinase is responsible for Abi-1-S323 phosphorylation. **(A-C)** DBTRG, H520, and SCC90 cells were treated with the CHK1 inhibitor Rabusertib **(A)**, the CHK2 inhibitor PV1019 **(B)**, or the NUAK1 inhibitor HTH-01-015 **(C)**, at the indicated concentrations for 24 hours and then harvested for immunoblotting analysis. Abi-1-S323 phosphorylation, CHK1 S296 phosphorylation, CHK2 S516 phosphorylation, and the downstream NUAK1 substrate MYPT1 S668 phosphorylation were analyzed to confirm target engagement and assess the effects of kinase inhibition on Abi-1 phosphorylation. **(D)** DBTRG, H520, and SCC90 cells were treated with 5 µM, 1 µM, 1 µM Rabusertib, respectively. Cells were harvested at indicated time point and analyzed by immunoblotting.

### CHK1 inhibition preferentially radiosensitizes tumors with low GTP-Rac1-Abi-1 activity

We previously identified that dephosphorylated Abi-1 promotes NHEJ. Because CHK1 is a canonical regulator of HR repair (45,46), the newly-defined ability of CHK1 to phosphorylate Abi-1 suggests that the Rac1-Abi-1 signaling axis may influence DSB repair pathway choice. We hypothesized that cells with high Rac1-Abi-1 signaling preferentially engage NHEJ, whereas cells with low Rac1-Abi-1 signaling rely more on CHK1-associated HR. Consistent with this hypothesis, H520 and SCC90 cells, which exhibited higher p-Abi-1-S323 levels, also showed higher basal CHK1 activity than H460 and Detroit 562 cells, which exhibited high Rac1 activity and low p-Abi-1-S323 (**Fig. 2**; **Fig. 7A** and **7D**). Following RT, NHEJ marker 53BP1 foci predominated in H460 and Detroit 562 cells, whereas HR marker RAD51 foci were more prominent in H520 and SCC90 cells (**Fig. 7B** and **7C**; **Fig. 7E** and **F**; Supplementary **Fig. S7A** and **S7B**). These findings support a model in which elevated Rac1-Abi-1 signaling favors NHEJ, whereas low Rac1-Abi-1 signaling is associated with increased CHK1 activity and preferential engagement of HR-mediated DSB repair.

**Figure 7.**
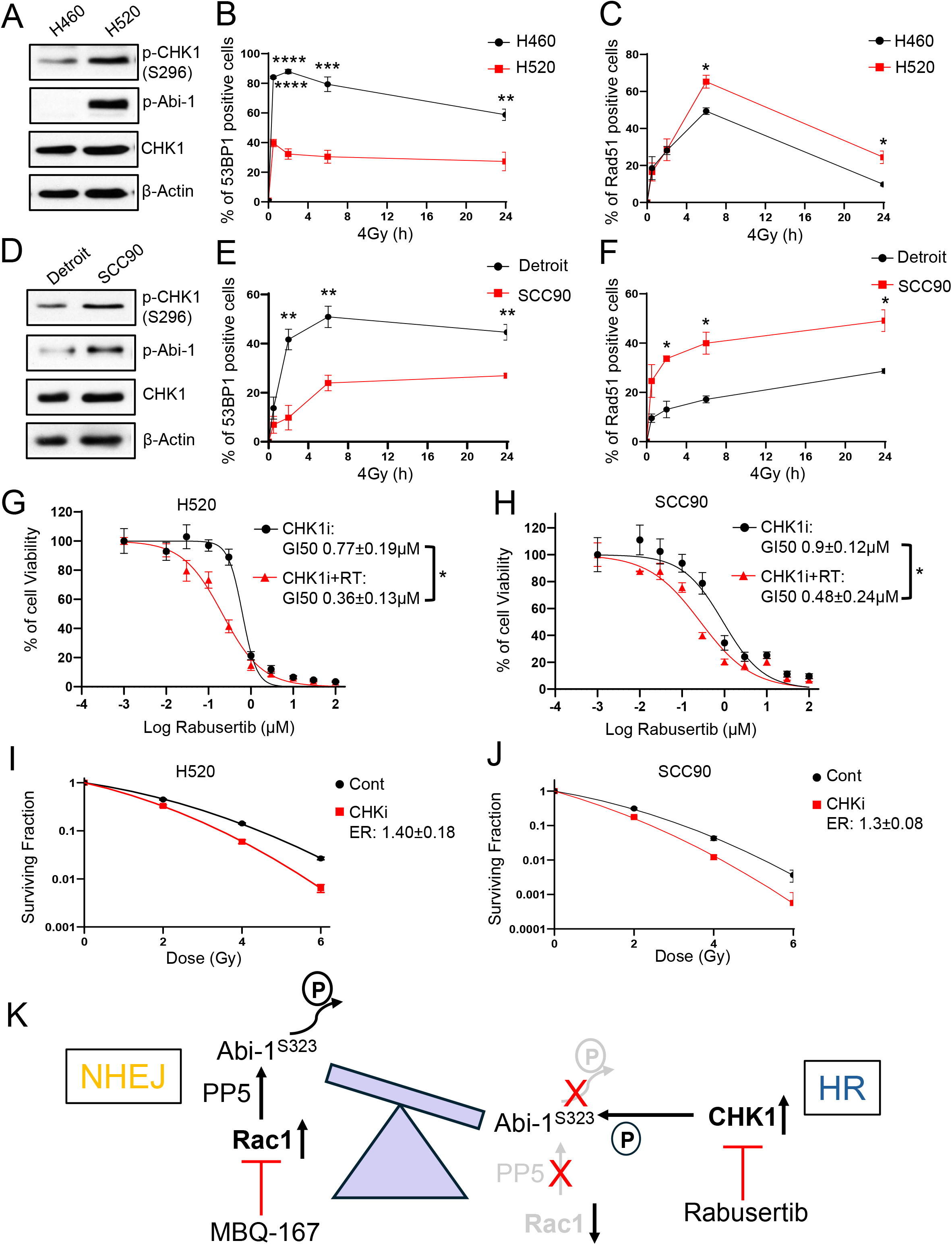
CHK1 inhibition preferentially enhances RT sensitivity in cancers with low Rac1-Abi-1 signaling activity and HR pathway dependency. **(A-F)** Cells were harvested for immunoblotting assay to determine the indicated protein levels **(A, D)**, or exposed to 4 Gy irradiation and harvested at 0.5, 2, 6, and 24 hours for immunofluorescence co-staining of 53BP1 **(B, E)** and RAD51 **(C, F)**. Foci thresholds were set at 10 for 53BP1 and 5 for Rad51. Representative immunoblots from three biological replicates are shown in **(A)** and **(D)**. For **(B)**, **(C)**, **(E)**, and **(F)**, *, p < 0.05; **, p < 0.01, ***, p < 0.001, ****, p < 0.0001 comparing the two groups at the corresponding time point. Data are presented as mean ± SEM from 3-4 independent biological replicates and were analyzed using a two-tailed *t* test. **(G-H)** H520 and SCC90 cells were treated with escalating concentrations of the CHK1 inhibitor Rabusertib alone or in combination with RT (4 Gy). Rabusertib was administered 2-4 hours before RT. Cell viability was assessed by CellTiter-Glo assay 5 days after irradiation, and GI₅₀ values were calculated. Data are presented as mean ± SEM from 3-4 independent biological replicates. *, p < 0.05; two-tailed *t* test. **(I-J)** Cells were treated overnight with the CHK1 inhibitor Rabusertib (1 µM), replated, and subjected to irradiation for clonogenic survival assays. Colonies were stained and counted 10-14 days after irradiation for H520, and 21 days after irradiation for SCC90 cells. ER (mean ± SD) from three biologic replicates is shown. **(K)** Schematic summary of the study.

We next investigated whether CHK1 inhibition preferentially radiosensitizes H520 and SCC90 cells, which exhibit low Rac1-Abi-1 signaling and high CHK1 activity. Long-term CellTiter-Glo assays showed that H520 and SCC90 cells were more sensitive to CHK1 inhibitor Rabusertib alone than H460 (GI₅₀: 0.77 ± 0.19 vs. 2.47 ± 0.21 μM; p = 0.0001; **Fig. 7G**; Supplementary **Fig. S7C**) and Detroit 562 cells (0.90 ± 0.12 vs. 1.55 ± 0.52 μM; p = 0.05; **Fig. 7H**; Supplementary **Fig. S7D**). Moreover, Rabusertib in combination with RT significantly enhanced growth inhibition in H520 (p = 0.01; **Fig. 7G**) and SCC90 (p = 0.04; **Fig. 7H**), but not H460 (p = 0.65; Supplementary **Fig. S7C**) or Detroit 562 (p = 0.45; Supplementary **Fig. S7D**) cells, compared with Rabusertib alone. Consistent with these findings, clonogenic survival assays showed minimal radiosensitization by Rabusertib in H460 (ER = 1.06 ± 0.07; Supplementary **Fig. S7E**) and Detroit 562 (ER = 1.07 ± 0.02; Supplementary **Fig. S7F**) cells, which exhibit elevated Rac1-Abi-1-NHEJ signaling and are preferentially sensitized by Rac1 inhibition (**Fig. 4**). In contrast, Rabusertib significantly enhanced RT sensitivity in H520 (ER = 1.40 ± 0.18; **Fig. 7I**) and SCC90 (ER = 1.30 ± 0.08; **Fig. 7J**) cells. Together, these findings demonstrate that Rac1-Abi-1 signaling status is associated with distinct DSB repair dependencies and suggest that tumors with low Rac1-Abi-1 signaling may be preferentially vulnerable to CHK1 inhibition, whereas those with elevated Rac1-Abi-1 signaling may benefit from Rac1-targeted radiosensitization.

## Discussion

In the present study, we identify p-Abi-1-S323 as a biomarker of RT response in patients and establish the Rac1-Abi-1 signaling axis as a previously unrecognized determinant of DNA repair pathway choice and therapeutic response across multiple cancer types. We demonstrate that loss of Abi-1-S323 phosphorylation predicts poorer outcomes in patients with GBM receiving RT, while bioinformatic analyses identify NSCLC and HNC as cancer types with high *RAC1* amplification. Consistent with these findings, loss of Abi-1-S323 phosphorylation before RT predicts poorer outcomes in patients with HNC. Our *in vitro* studies further demonstrate that elevated Rac1 activity, accompanied by Abi-1-S323 dephosphorylation, promotes efficient DSB repair and intrinsic radioresistance in NSCLC and HNC models, whereas genetic or pharmacological inhibition of the Rac1-Abi-1 axis impairs DNA repair and enhances RT sensitivity both *in vitro* and *in vivo*. We identify CHK1 as the kinase responsible for Abi-1-S323 phosphorylation and demonstrate that the Rac1-Abi-1 axis is associated with distinct functional DNA repair states. Tumors with high Rac1 activity and low Abi-1 phosphorylation exhibit an NHEJ-dominant state associated with reduced CHK1 activity and are preferentially sensitized to Rac1 inhibition but relatively resistant to CHK1 inhibition. In contrast, tumors with low Rac1 activity and high Abi-1 phosphorylation exhibit greater CHK1 activity and an HR-dependent repair state, rendering them preferentially sensitive to CHK1 inhibition (**Fig. 7K**).

Our findings suggest that Abi-1-S323 phosphorylation may represent a functional biomarker of tumor response to genotoxic RT in human cancers. Despite decades of investigation, clinically applicable biomarkers that reliably predict RT response remain limited. Several biomarkers have been associated with RT response in specific tumor types, including HPV positivity in HNC(37,47), and MGMT promoter methylation in GBM(2). Likewise, BRCA1/2 mutations and other forms of HR deficiency (HRD) confer impaired DSB repair and increased sensitivity to DNA-damaging therapies(48). However, these biomarkers are largely applied to specific molecular subsets of tumors. More broadly relevant factors, including TP53 mutation(35) and tumor hypoxia(49), have also been associated with radioresistance, but their predictive utility is limited by tumor heterogeneity or inconsistent clinical performance. In contrast, Abi-1-S323 dephosphorylation reflects a functional DNA repair state characterized by Rac1-dependent NHEJ activity. Because Rac1 activation can be driven by dysregulated GTP metabolism, a fundamental metabolic process frequently altered in cancer, this pathway may provide a biologically integrated measure of tumor DNA repair capacity. Notably, the association between p-Abi-1 and RT response was observed across multiple cancer types, including brain(22), lung, and head and neck cancers (this study), and across experimental systems ranging from cell and animal models to patient tumors.

The association of the Rac1-Abi-1 axis with distinct DNA repair states provides a rationale for context-dependent use of DDR inhibitors. CHK1 inhibitors have shown potent radiosensitizing activity in preclinical models by disrupting RT-induced checkpoint activation, replication fork protection, and DNA repair (50,51), and have been evaluated in clinical trials (NCT02203513; NCT03414047). However, clinical responses have been variable, underscoring the need for biomarkers that identify tumors with functional CHK1 dependency. Our findings demonstrate that Rac1-Abi-1 signaling status is associated with differential sensitivity to CHK1 inhibition. Tumors with low Rac1 activity and high Abi-1 phosphorylation exhibit greater CHK1 activity and an HR-dependent repair state and are consequently more sensitive to CHK1 inhibition. These findings suggest that functional biomarkers defining tumor DNA repair states may enable more precise identification of tumors dependent on CHK1-mediated repair and more likely to benefit from CHK1 inhibitor-based radiosensitization. Consistent with this model, although p-Abi-1-positive HNC tumors were associated with more favorable initial clinical outcomes (**Fig. 1E** and **1F**), a subset subsequently developed recurrence. This observation raises the possibility that p-Abi-1-positive tumors may represent a clinically relevant population for future evaluation of CHK1 inhibitor-based radiosensitization strategies.

Our findings provide a strong rationale for advancing MBQ-167 in combination with RT toward clinical evaluation in patients with tumors exhibiting high Rac1-Abi-1 signaling activity. Although several Rac1 inhibitors have demonstrated anti-tumor or radiosensitizing effects in preclinical models (52,53), they remain in early-stage development. MBQ-167, a dual Rac1/CDC42 inhibitor, has demonstrated promising antitumor and antimetastatic activity in preclinical studies (31,38) and is currently undergoing clinical evaluation (NCT06075810). Our recent work established that GTP-Rac1-Abi-1 signaling promotes NHEJ and RT resistance in GBM and that Rac1 inhibition with MBQ-167 enhances RT sensitivity in orthotopic patient-derived xenograft models(22). The present study demonstrates effective tumor drug delivery, target engagement, and potent radiosensitizing activity of MBQ-167 across multiple cancer models *in vivo* (**Fig. 5**). Importantly, MBQ-167 did not enhance RT-induced cytotoxicity in normal cells (Supplementary **Fig. S4A** and **S4B**), suggesting a potential therapeutic window that may reflect greater dependence of tumor cells on Rac1-Abi-1 signaling for DNA repair. Consistent with this possibility, Rac1 activity has been reported to increase preferentially in tumor cells following RT (52), potentially creating a greater dependence on Rac1-mediated DNA repair.

Our study also raises several important questions and limitations. First, the mechanisms underlying the differential dependence on Rac1-Abi-1 signaling between cancer and normal cells remain unclear. Second, whether tumors with low Rac1 activity and high Abi-1 phosphorylation engage DNA repair pathways beyond HR warrants further investigation. Third, given the physiological roles of Rac1 and Abi-1 in normal cells, strategies that selectively target their DNA repair function, including direct modulation of Abi-1 phosphorylation, may offer a broader therapeutic window than systemic Rac1 inhibition. Finally, the prognostic and predictive value of p-Abi-1 requires validation in larger, diverse cohorts and prospective studies. In conclusion, our study establishes the Rac1-Abi-1 axis as a regulator of tumor DNA repair states and identifies Abi-1 S323 phosphorylation as a potential therapeutic biomarker.

## Authors’ Contributions

**Y Huang, E McCulla, J Park, and A Yang**: data curation, formal analysis, validation, investigation, visualization, methodology. Y Huang: writing original draft. **T Jho, A. Lin, J Xu, L Hess**: Validation, investigation, methodology. **K. Wilder-Romans**: Investigation, methodology. **J. Li**: Resources. **N Liang, Z Zhu, D Kothari, K Jin, and S Kim**: Investigation, methodology. **S Gedara, B. Wen and D Sun**: Methodology. **M. Vinco and S.P. Ferris**: Investigation, methodology. **M.A. Morgan and T.S. Lawrence:** Writing–review and editing. **Y.M. Shah:** Resources, writing–review and editing. **M. Yu:** data analysis. **ME. Heft Neal**: Resources, methodology, writing–review and editing. **J.C. Brenner:** Resource. **S Dharmawardhane and J.F. Rodriguez-Orengo:** Resource**. D.R. Wahl:** resources, writing–review and editing. **W Zhou**: Conceptualization, resources, data curation, formal analysis, supervision, funding acquisition, visualization, methodology, project administration, writing–review and editing.

## Acknowledgments

Animal management, technical and experimental support services were provided by the University of Michigan Unit for Laboratory Animal Medicine (ULAM). E. McCulla was supported by the NCI Training Grant T32-CA009676, 1F31CA31406801. D.R. Wahl was supported by the NCI (R37CA258346), and the National Institute of Neurological Disorders and Stroke (R01NS129123). W. Zhou was supported by the UMMS Pandemic Research Recovery Program (U083054), Rogel Cancer Center Discovery Award and Michigan Radiosensitization SPORE Career Enhancement Program (5-P50-CA-269022-02). M. A. Morgan was supported by NCI (P50CA269022). L. D. Hess was supported by the NSF GRFP (DGE2241144). Research reported in this publication was partly supported by the National Cancer Institute of the National Institutes of Health under Award Number P30CA046592 through the use of the Rogel Cancer Center Experimental Irradiation Shared Resource and Cancer Data Science Shared Resource. The content is solely the responsibility of the authors and does not necessarily represent the official views of the National Institutes of Health. Research reported in this publication was supported by the National Cancer Institute of the National Institutes of Health under award number P30CA046592 for pharmacokinetic study. Research reported in this publication was supported by Rogel Comprehensive Cancer Center (P30CA046592) through the Experimental Irradiation Shared Resource.

## Notes

### Competing Interest Statement

D.R.W. has consulted for Agios Pharmaceuticals, Admare Pharmaceuticals, and Bruker and Innocrin Pharmaceuticals. D.R.W. is an inventor on patents pertaining to the treatment of patients with brain tumors (U.S. Provisional Patent Application 62/744,342, U.S. Provisional Patent Applicant 62/724,337).
Meredith A. Morgan has received funds and served on an advisory committee for AstraZeneca.
The other authors declare no competing interests.

