## Supplemental Figures for "Rac1 and CHK1 Converge on Abi-1 to Regulate DNA Repair Dependency and Treatment Response in Human Cancers"

*Huang et al.*, Figure S1

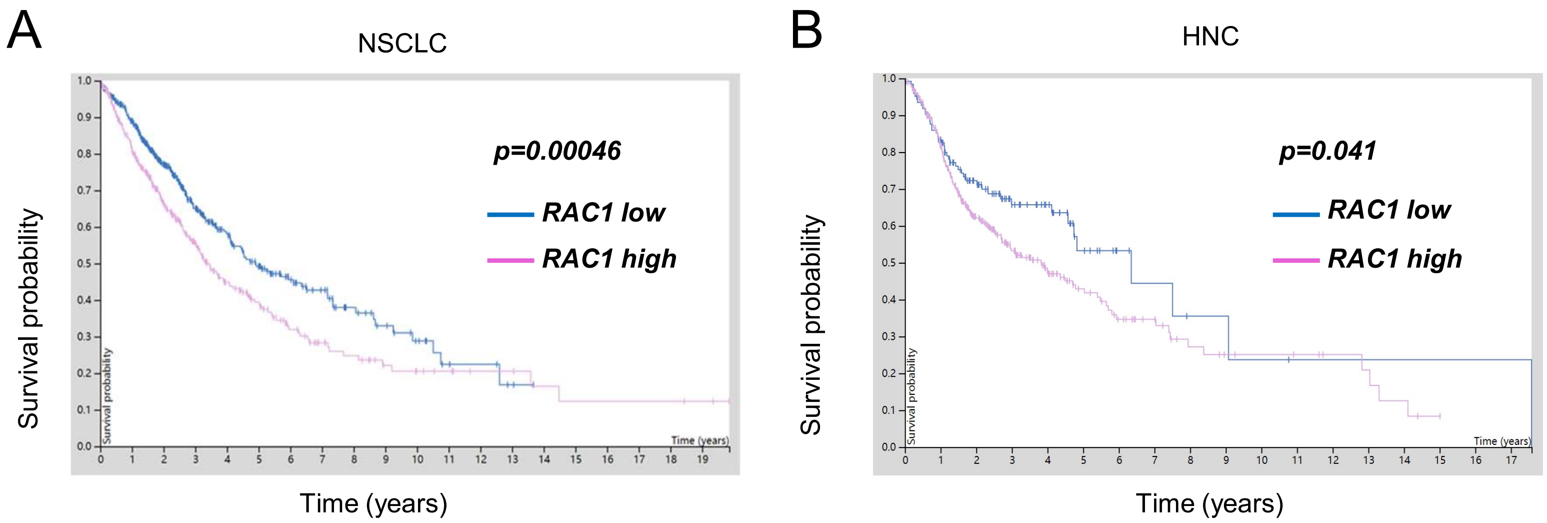

*Huang et al.*, Figure S2

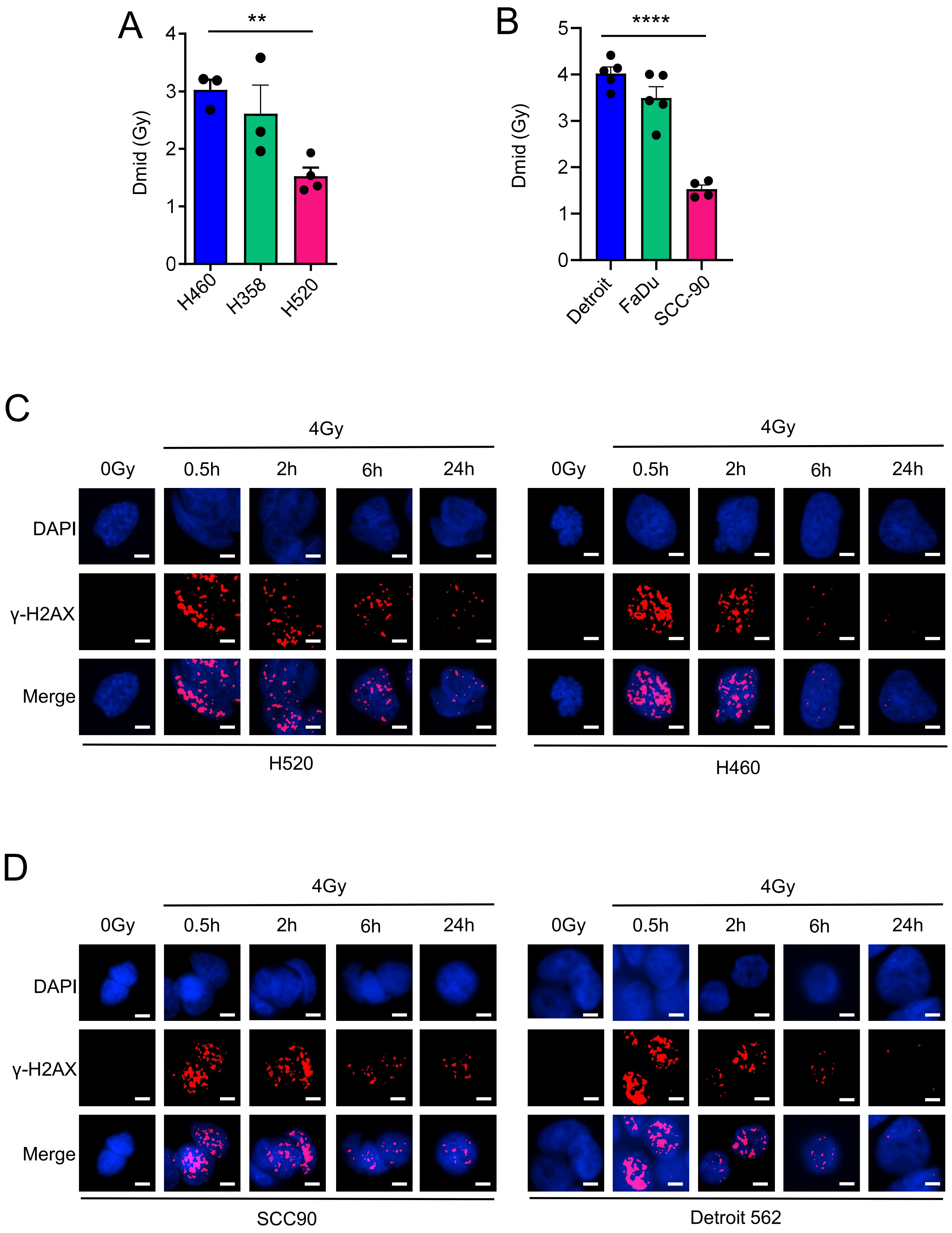

*Huang et al.*, Figure S3

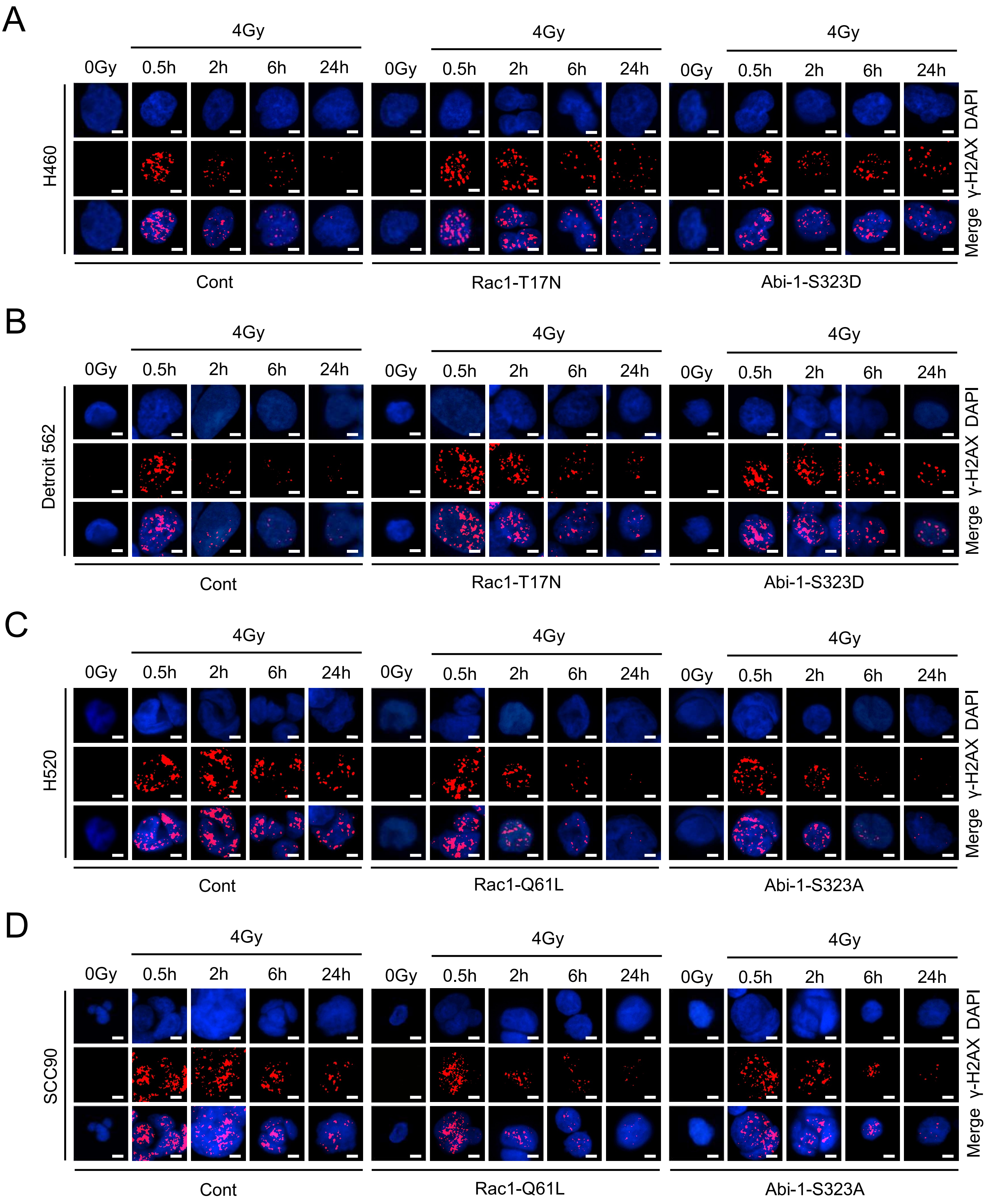

*Huang et al.,* Figure S4

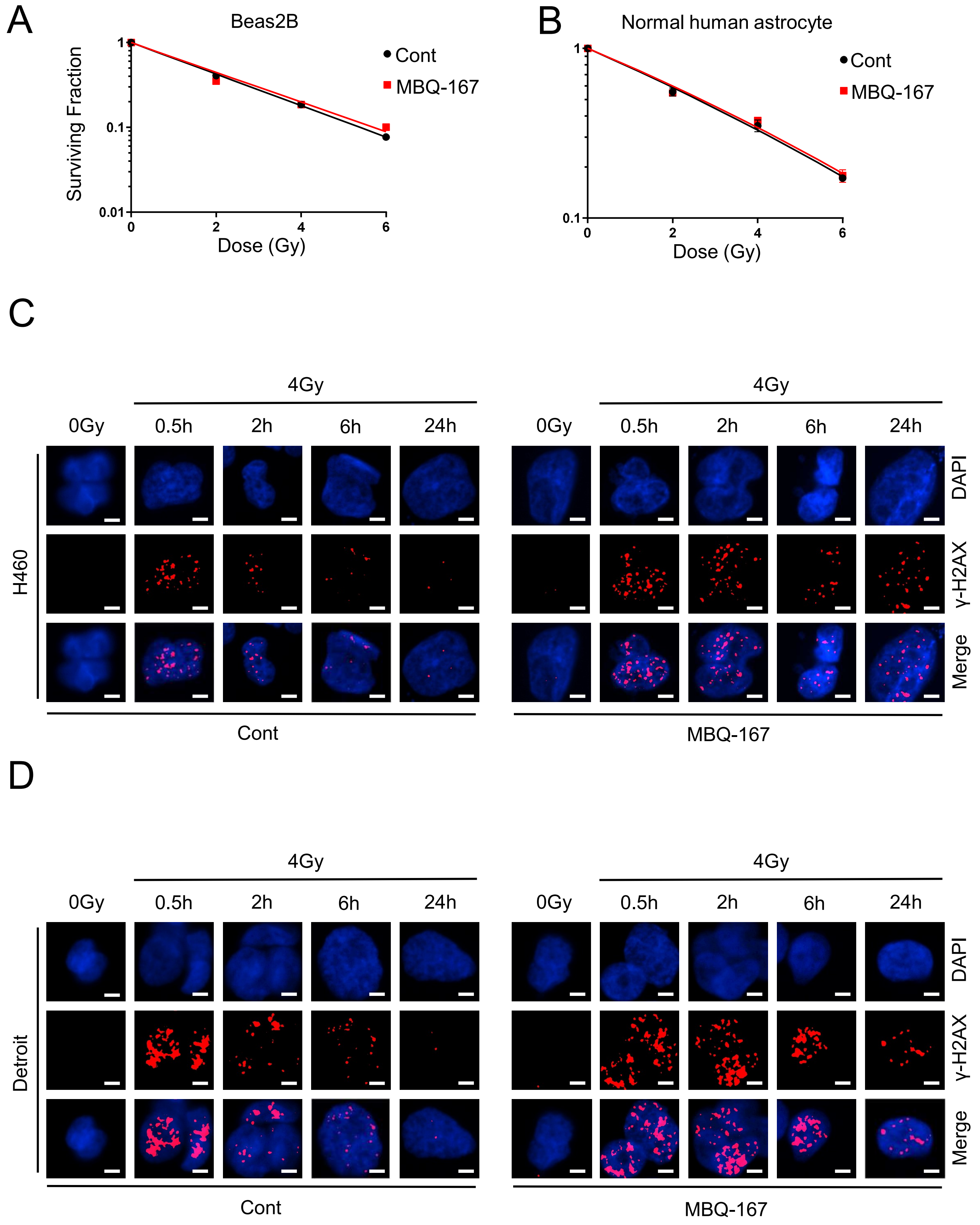

*Huang et al.,* Figure S5

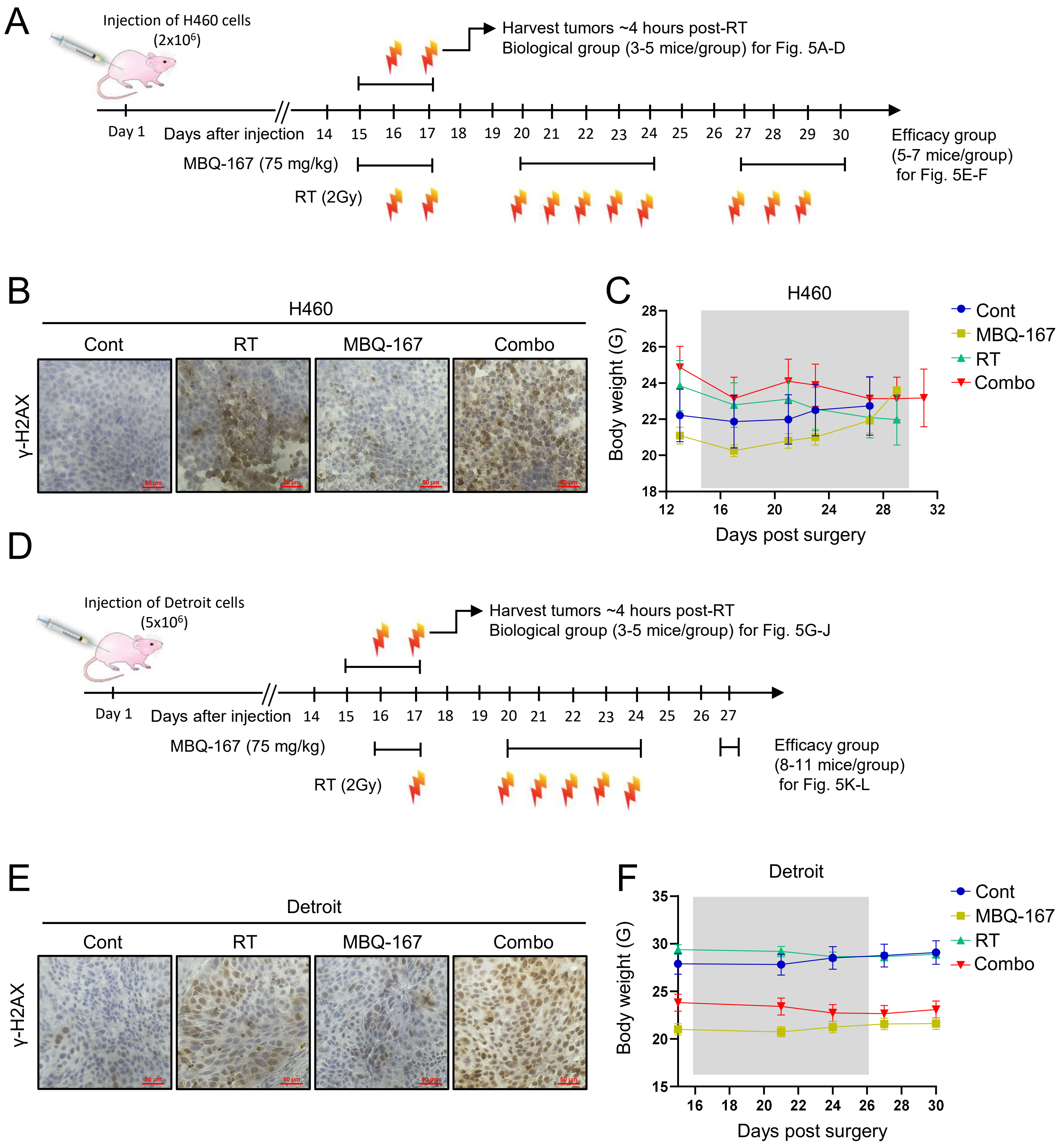

*Huang et al.*, Figure S6

**A** Potential kinases that phosphorylate Abi-1 S323

| Kinase Name | Percentile % | Ranking |
|---|---|---|
| CHK1 | 99.87 | 1 |
| CHK2 | 98.89 | 2 |
| NUAK1 | 98.87 | 3 |
| SBK | 98.32 | 4 |
| RIPK1 | 98.31 | 5 |
| PKN1 | 98.25 | 6 |
| DLK | 98.16 | 7 |
| CAMK4 | 97.82 | 8 |
| PKN2 | 97.68 | 9 |
| BCKDK | 97.66 | 10 |

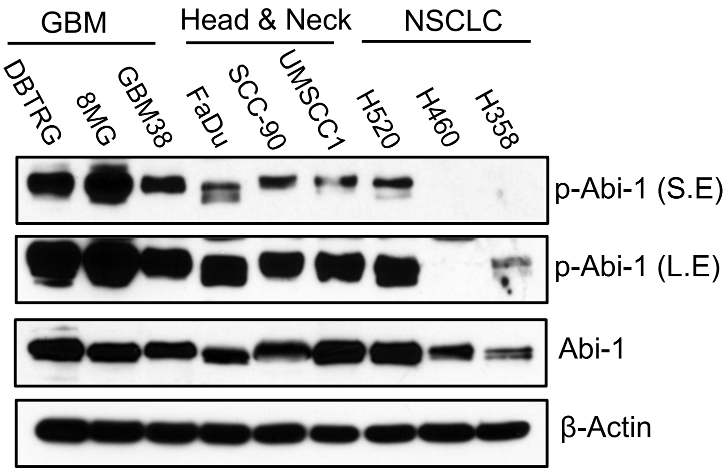


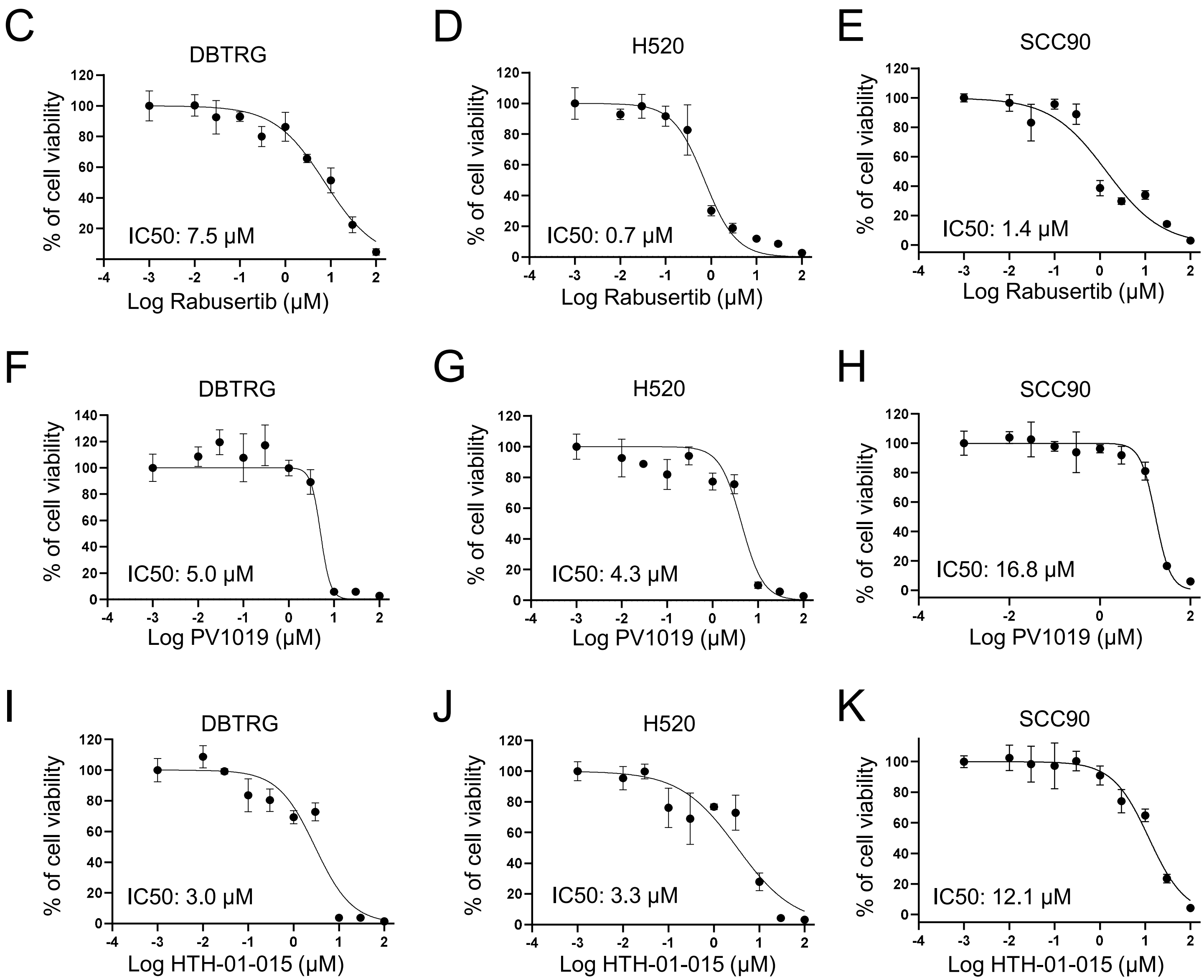

Huang et al., Figure S7

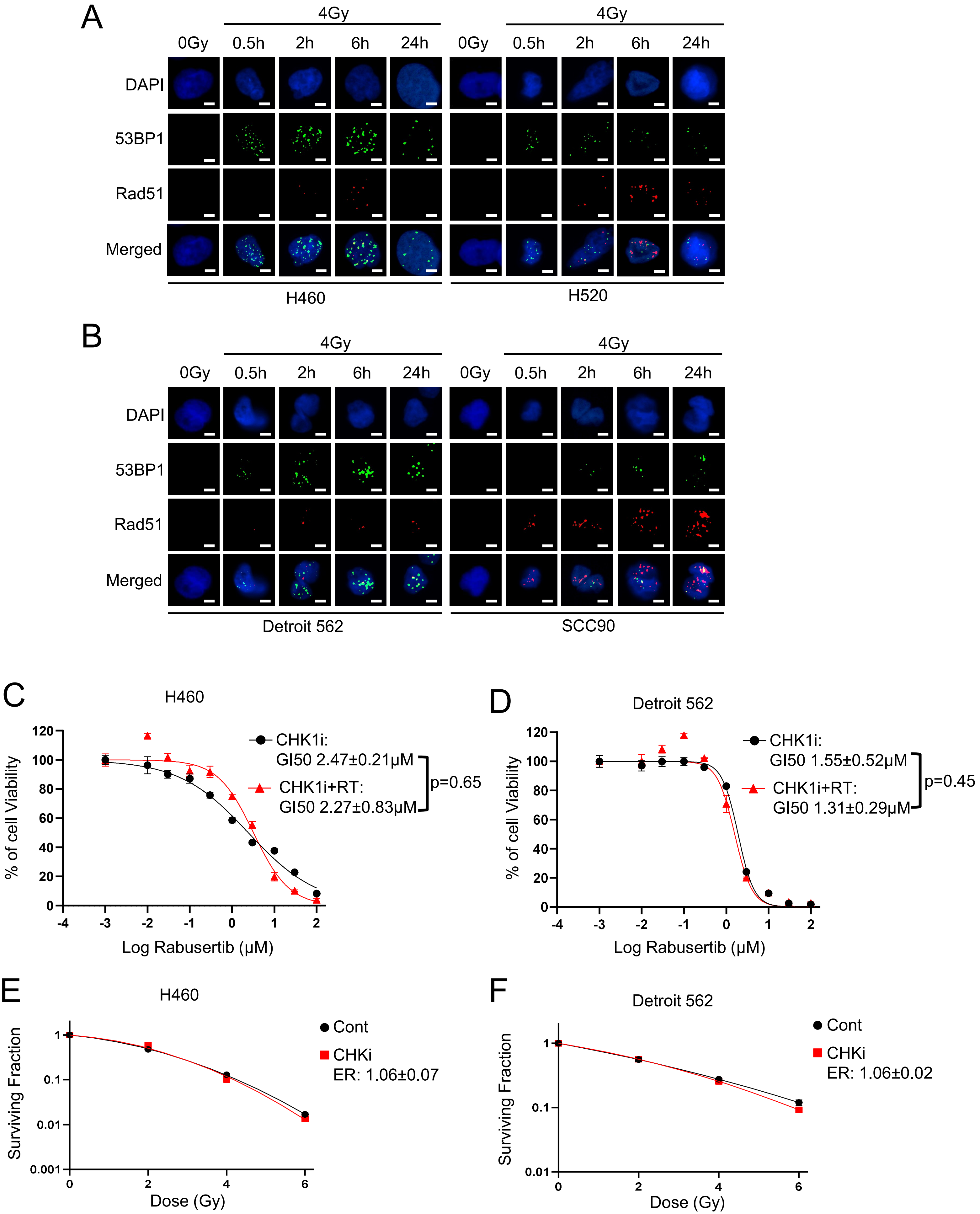
