## Supplemental Information for "Rac1 and CHK1 Converge on Abi-1 to Regulate DNA Repair Dependency and Treatment Response in Human Cancers"

[1]Department of Radiation Oncology, University of Michigan, Ann Arbor, Michigan.
[2]Rogel Cancer Center, University of Michigan Medical School, Ann Arbor, Michigan.
[3]Doctoral Program in Cancer Biology, University of Michigan Medical School, Ann Arbor, Michigan
[4]Cell Signaling Technology, Inc., Danvers, Massachusetts.
[5]Department of Pharmaceutical Sciences, College of Pharmacy, University of Michigan, Ann Arbor, Michigan.
[6]Department of Pathology, Tissue & Molecular Pathology Division, University of Michigan, Ann Arbor, Michigan.
[7]Department of Pathology, Division of Neuropathology, University of Michigan, Ann Arbor, Michigan.
[8]Department of Molecular and Integrative Physiology, University of Michigan, Ann Arbor, Michigan
[9]School of Public Health and Department of Biostatistics, University of Michigan, Ann Arbor, Michigan
[10]Department of Otolaryngology-Head and Neck Surgery, University of Michigan, Ann Arbor, Michigan
[11]Department of Pharmacology, University of Michigan, Ann Arbor, Michigan.
[12]MBQ Pharma, Inc. Puerto Rico
[13]Department of Biochemistry, School of Medicine, University of Puerto Rico
[14]Department of Neurosurgery, University of Michigan, Ann Arbor, Michigan
*These authors contributed equally.

**Running title**: RAC1 and CHK1 Determine DNA Repair Dependency via Abi-1

**Corresponding author:**
Weihua Zhou
Department of Radiation Oncology,
University of Michigan,
Ann Arbor, MI, USA


### SUPPLEMENTAL INFORMATION

#### Supplemental Figure Legends

**Figure. S1. High *RAC1* mRNA levels predict worse clinical outcomes in NSCLC and HNC.** **(A-B)** *RAC1* mRNA expression and clinical information were obtained from The Cancer Genome Atlas (TCGA). Kaplan–Meier analyses of overall survival (OS) in NSCLC and HNC cohorts stratified by *RAC1* expression are shown.

**Figure. S2. High Rac1-Abi-1 signaling activity correlates with radioresistance in NSCLC and HNC.**

**(A-B)** NSCLC cells (H358, H520, and H460) and HNC cells (Detroit 562, FaDu, and SCC90) were exposed to escalating doses of RT and analyzed by clonogenic survival assay. The averaged Dmid, defined as the mean inactivating radiation dose and calculated as the area under the clonogenic survival curve, is shown based on 3–5 independent biological replicates (Fig. S2A corresponding to Fig. 2D and Fig. S2B corresponding to Fig. 2H). **, $p < 0.01$; ****, $p < 0.001$; mean ± SEM (two paired t test). **(C-D)** H520 and H460 cells **(C)**, as well as SCC90 and Detroit 562 cells **(D)** were irradiated with 4 Gy and fixed 0.5, 2, 6, 24 hours later for immunofluorescence staining of γ-H2AX. Representative IF images of γ-H2AX foci staining in H520 and H460 (Fig. S2C corresponding to Fig. 2I) as well as SCC90 and Detroit 562 (Fig. S2D corresponding to Fig. 2J) were shown. Scale bar, 5 μm.

**Figure S3. Genetic modulation of Rac1 activity and Abi-1 phosphorylation regulates DSB repair and radioresistance in NSCLC and HNC.**

**(A-B)** H460 and Detroit 562 (Detroit) cells were transiently transfected with myc-tagged Rac1-T17N or Flag-tagged Abi-1-S323D, followed by r-H2AX IF foci staining 0.5, 2, 6, 24 hours after RT. Representative IF images of γ-H2AX foci staining in H460 (corresponding to Fig. 3C) and Detroit 562 (corresponding to Fig. 3F). **(C-D)** H520 and SCC90 cells were transiently transfected with myc-tagged Rac1-Q61L or Flag-tagged Abi-1-S323A, followed by r-H2AX IF foci staining 0.5, 2, 6, 24 hours after RT. Representative IF images of γ-H2AX foci staining in H520 (corresponding to Fig. 3I) and SCC90 (corresponding to Fig. 3L) were shown. Scale bar: 5 μm.

**Figure S4. Pharmacological inhibition of Rac1-Abi-1 signaling enhances radiosensitivity by suppressing DNA DSB repair.**

**(A-B)** Normal bronchial epithelial cells (Beas2B) and normal human astrocytes were treated with MBQ-167 (100 nM) overnight and then replated for clonogenic survival assays following RT. Representative clonogenic survival curves from one independent biological replicate are shown. **(C-D)** H460 and Detroit 562 (Detroit) cells were treated with MBQ-167 overnight, replated, and fixed for γ-H2AX immunofluorescence staining 0.5, 2, 6, 24 hours after irradiation (4 Gy). Representative images of γ-H2AX foci in H460 and Detroit 562 cells are shown, corresponding to Fig. 4E and 4F, respectively. Scale bar: 5 μm.

**Figure S5. Pharmacological inhibition of Rac1-Abi-1 signaling enhances radiosensitivity in xenograft tumor models.**

**(A)** Schematic timeline of the H460 flank xenograft model. H460 cells ($2 \times 10^6$) were injected subcutaneously into the flanks of mice, and tumors were allowed to establish. Mice were then randomized into treatment groups as described in the Methods. MBQ-167 (75 mg/kg) was administered by oral gavage once daily, 2 h before RT (2 Gy/fraction) on weekdays. For pharmacodynamic analyses, a subset of mice (3-5 per group; upper timeline) received three doses of MBQ-167 and two fractions of RT, and tumors were harvested at ~4 h after the second RT fraction. A separate cohort of mice (n = 6-7 per group; lower timeline) received the full treatment regimen (12 doses of MBQ-167 and 10 fractions of RT) to evaluate therapeutic efficacy. **(B)** Representative IHC staining of γ-H2AX in H460 xenograft tumors harvested at ~4 h after the second RT fraction (corresponding to Fig. 5D). Scale bar: 50 μm. **(C)** Body weight curves of mice bearing H460 xenografts. Body weight was monitored throughout the treatment period (gray shaded area). **(D)** Schematic timeline of the Detroit 562 (Detroit) flank xenograft model. Detroit cells ($5 \times 10^6$) were injected subcutaneously into the flanks of mice, and tumors were allowed to establish. A subset of mice (3-5 per group; upper timeline) received three doses of MBQ-167 and two fractions of RT, and tumors were harvested 4 h after the second RT fraction. A separate cohort of mice (n = 8-11 per group; lower timeline) received the full treatment regimen (8 doses of MBQ-167 and 6 fractions of RT) to evaluate therapeutic efficacy. **(E)** Representative IHC staining of γ-H2AX in Detroit 562 xenograft tumors (corresponding to Fig. 5J). Scale bar: 50 μm. **(F)** Body weight curves of mice bearing Detroit 562 xenografts. Body weight was monitored throughout the treatment period (gray shaded area).

**Figure S6. CHK1 kinase is responsible for Abi-1 S323 phosphorylation.**

**(A)** Summary of the top 10 candidate kinases predicted to be responsible for Abi-1 phosphorylation at Ser323. **(B)** Cell pellets from GBM, NSCLC, and HNC cell lines were collected and subjected to immunoblotting using the indicated antibodies. **(C-K)** DBTRG, H520, and SCC90 cells were treated with escalating concentrations of the CHK1 inhibitor Rabusertib, the CHK2 inhibitor PV1019, or the NUAK1 inhibitor HTH-01-015 for 24 hours, followed by CellTiter-Glo assays to determine the $GI_{50}$ values of each inhibitor.

**Figure S7. CHK1 inhibition preferentially enhances RT sensitivity in cancers with low Rac1-Abi-1 signaling activity and HR pathway dependency.**

**(A-B)** H460 and H520 cells **(A)**, as well as Detroit 562 and SCC90 cells **(B)**, were exposed to 4 Gy irradiation and harvested at 0.5, 2, 6, and 24 hours after irradiation for immunofluorescence (IF) staining of 53BP1 and RAD51 foci. Representative immunofluorescence images of 53BP1 and RAD51 foci in H460 and H520 cells (corresponding to Fig. 7B and 7C) and Detroit and SCC90 cells (corresponding to Fig. 7E and 7F) are shown. Scale bar: 5 μm. **(C-D)** H460 and Detroit 562, cells were treated with escalating concentrations of the CHK1 inhibitor Rabusertib alone or in combination with RT (4 Gy). Rabusertib was administered 2-4 hours before RT. Cell viability was assessed by CellTiter-Glo assay 5 days after irradiation, and $GI_{50}$ values were calculated. Data are presented as mean ± SEM from 4 independent biological replicates; two-tailed *t* test. **(E-F)** Cells were treated overnight with the CHK1 inhibitor Rabusertib (1 μM), replated, and subjected to irradiation for clonogenic survival assays. Colonies were stained and counted 10–14 days after irradiation. ER (mean ± SD) from three biologic replicates is shown.
